# Capture at the single cell level of metabolic modules distinguishing aggressive and indolent glioblastoma cells

**DOI:** 10.1101/755611

**Authors:** Mirca S. Saurty, Léa Bellenger, Elias A. El-Habr, Virgile Delaunay, Hervé Chneiweiss, Christophe Antoniewski, Ghislaine Morvan-Dubois, Marie-Pierre Junier

**Affiliations:** Sorbonne Université, CNRS UMR8246, Inserm U1130, Neuroscience Paris Seine-IBPS, Team glial plasticity and neurooncology, Paris, France; ARTbio Bioinformatics Analysis Facility, Sorbonne Université, CNRS, Institut de Biologie Paris Seine, 75005 Paris, France

**Keywords:** Glioma, computational modeling, scRNA-seq, PUFA, elongase

## Abstract

Glioblastoma cell ability to adapt their functioning to microenvironment changes is a source of the extensive intra-tumor heterogeneity characteristic of this devastating malignant brain tumor. A systemic view of the metabolic pathways underlying glioblastoma cell functioning states is lacking. We analyzed public single cell RNA-sequencing data from glioblastoma surgical resections, which offer the closest available view of tumor cell heterogeneity as encountered at the time of patients’ diagnosis. Unsupervised analyses revealed that information dispersed throughout the cell transcript repertoires encoded the identity of each tumor and masked information related to cell functioning states. Data reduction based on an experimentally-defined signature of transcription factors overcame this hurdle. It allowed cell grouping according to their tumorigenic potential, regardless of their tumor of origin. The approach relevance was validated using an independent dataset of glioblastoma tissue transcriptomes, patient-derived cell lines and orthotopic xenografts. Overexpression of genes coding for amino acid and lipid metabolism enzymes involved in anti-oxidative, energetic and cell membrane processes characterized cells with high tumorigenic potential. Modeling of their expression network highlighted the very long chain polyunsaturated fatty acid synthesis pathway at the core of the network. Expression of its most downstream enzymatic component, ELOVL2, was associated with worsened patient survival, and required for cell tumorigenic properties in vivo. Our results demonstrate the power of signature-driven analyses of single cell transcriptomes to obtain an integrated view of metabolic pathways at play within the heterogeneous cell landscape of patient tumors.

## Introduction

Glioblastoma (GBM), the most common form of malignant brain tumors in adults, are characterized by extensive cell heterogeneity which results from irreversible processes - clonal selection of distinct mutations and differentiation of cancer stem cells - but also from the cells’ ability to adapt their functioning to variations in their environment and to therapies [2, 13, 32]. As a result, cancer cells coexist within GBM micro-territories in various functioning states, with respect to stem-like, proliferation, metabolic, migration, pro-angiogenic, drug resistance, or tumor-initiating (i.e. tumorigenic) capacities [9, 16, 47, 66]. Such heterogeneity in cell functioning defies therapeutic targeting.

The changes in cell functioning state are accompanied by variations in cell metabolic activities. These variations are essential for GBM cells to exploit different sources of nutrients such as glucose, glutamine or acetate, and thereby cope with changes in oxygen and nutrient availabilities that occur throughout tumor development [45, 46, 49]. The significance of these metabolic variations for the cell behavior may extend beyond a passive response to environmental signals, as recent evidence support a role for metabolism as a driver of changes in cell functional status. Flavahan and colleagues demonstrated that up-regulation of the high-affinity glucose transporter GLUT3 promotes acquisition by GBM cells of tumorigenic properties [23]. Conversely, we found that decreased activity of the mitochondrial enzyme SSADH triggers GBM cell conversion into a less aggressive functioning state, by coupling enhanced levels of the GABA by-product GHB to altered epigenetic regulations [18]. These metabolic variations have been found to take place within the patient tumors, and to be coherently linked with relevant phenotypic markers [18], or patients’ clinical course [12, 23]. Metabolism is also emerging as a player in GBM therapeutic resistance, as exemplified by escape from the anti-angiogenic Bevacizumab treatment. This escape has been linked to an increase in glycolysis and its uncoupling from oxidative phosphorylation in favor of lactate production in in vivo GBM models as well as in patients [19]. Metabolic enzymes are at the core of the molecular pathways controlling cell functioning states. Correcting their deregulation is therefore expected to be efficient to prevent acquisition and maintenance of aggressive cell functioning states shared by cell subpopulations in all GBM, regardless of their genomic specificities. Exploitation of metabolic targeting for therapies demands therefore to identify the metabolic pathways at play within the patient tumors in link with the heterogeneity of cell functioning states observed in GBM. Here, we used publicly available GBM single cell RNA-sequencing (scRNA-seq) data from four patients with EGFR amplification [14] for identifying metabolic pathways prevailing in GBM cell subpopulations in their most aggressive functioning state (Fig. 1A).

**Fig. 1.**
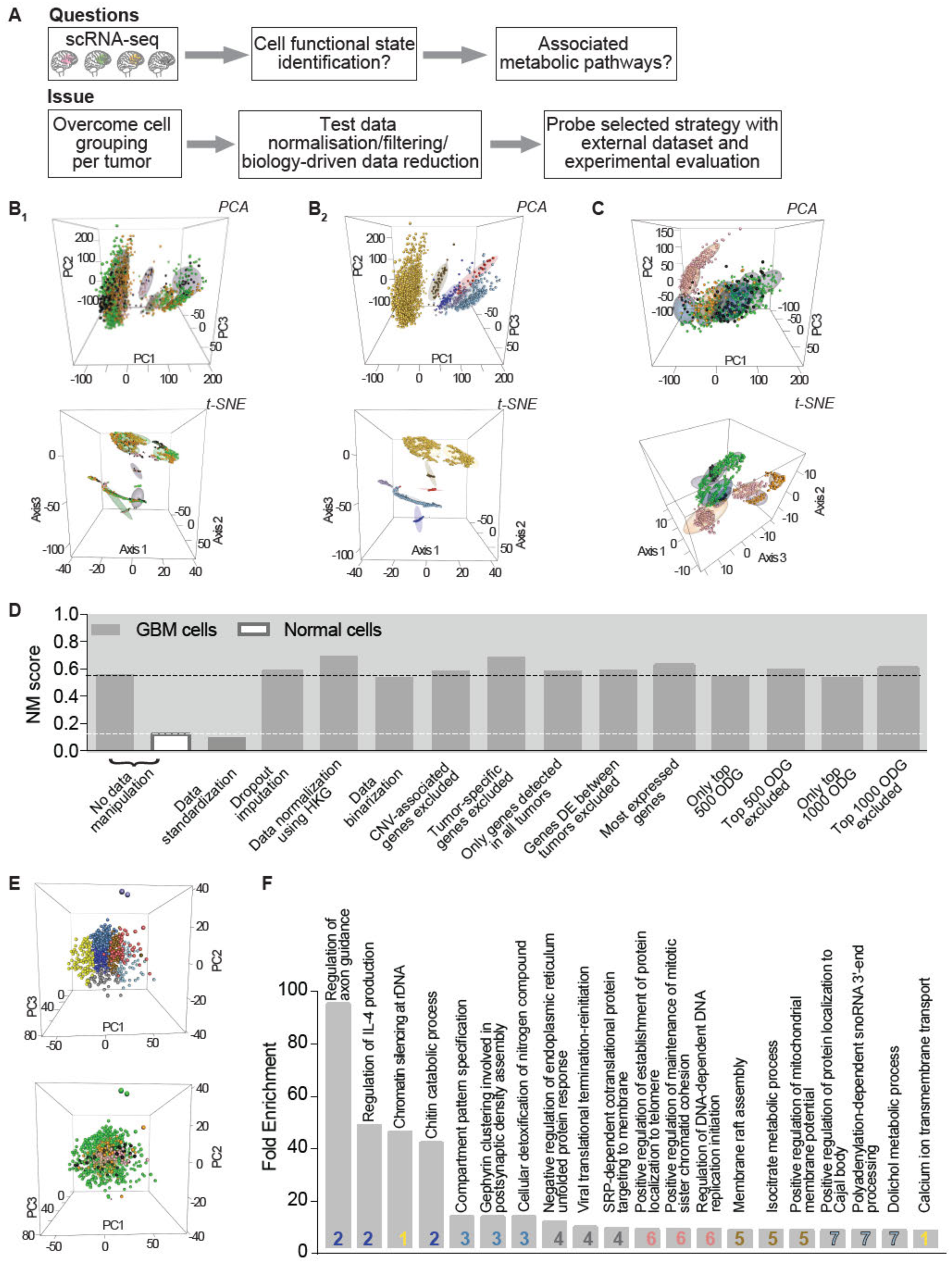
Spontaneous grouping of cancer cells by tumor of origin following unsupervised analysis. **a.** Schematic outline of the analytical and experimental strategy. **b-c.** Normal cells group independently from tumor of origin whereas cancer cell group by their tumor of origin. PCA and tSNE visualizations in upper and lower panels, respectively. Each dot represents a cell. b_1_ and c: cells colored by tumor of origin (pink, green, orange, black for GBM1, 2, 4 and 6, respectively). Ellipses delineate cell clusters identified upon unsupervised analysis. b_2_: cells colored by normal cell type identity (purple: astrocytes; blue: oligodendrocytes; light blue: oligodendrocyte precursor cells; red: neurons; gold: myeloid cells; brown: vascular cells). Ellipses delineate normal cell type identities as previously described [14]. **d.** Impact of data treatment on the dependence of cell clustering to tumors. NMI: Normalized Mutual Information score. Black and white dotted lines correspond to the reference NMI scores of grouping analyses performed with all genes detected in GBM and normal cells, respectively. Note that NMI scores of GBM cell grouping remain constant, regardless of the mode of data normalization or filtering (PCA plots in Fig. S2). Only data standardization reduces the NMI score to a value similar to that obtained when analyzing normal cells. **e.** Unsupervised analysis of data standardized by tumor results in clusters mixing cells from different tumors. PCA plots with highlight of the 7 clusters identified (top panel) or of the tumor from which the cells derive (bottom panel: pink, green, orange, black for GBM1, 2, 4 and 6, respectively). **f.** Gene ontology analysis of the genes describing each of the clusters highlight a variety of biological processes, not linkable to specific functioning states. DAVID toolkit. Corresponding cluster number is indicated (colored as cluster colors in the e upper panel).

Transcriptomes obtained by scRNA-seq are endowed with the potential to deliver information on a cell functioning state and its underlying molecular networks. The current analytic methods of scRNA-seq from different tumors result in the predominant grouping of cancer cells according to the tumor from which the cells are isolated (hereafter designed as tumor of origin), whereas normal cells present in the tumor are grouped according to their lineage subtype (e.g. neural, immune, vascular), regardless of their tumor of origin [14, 54, 61]. Characterizing the source of this specific influence of the tumor of origin on cancer cell grouping led to the development of a data reduction approach based on a molecular signature for identifying cell functioning states. Combining analyses of GBM unicellular and tissue transcriptomes with experimental assays, we highlighted a combination of metabolic pathways prevailing in cells with high tumorigenic potential. The analytical method developed is provided.

## Material and Methods

All figures were prepared using Adobe Illustrator (Adobe Systems). All bioinformatics analyses were performed using the R software version 3.5.0 (https://cran.r-project.org/). All resources and materials, R packages, corresponding websites and references are listed in Table S1. Detailed methods are provided in Supplementary Information 1.

### Computational analyses

Single cell transcriptomes of 1033 GBM and 2417 normal cells from four patients were used [14]. This dataset distinguishes cancer from normal cells on the basis of chromosome copy number variation (CNV) profiling, and further distinguishes normal cells according to their neural or immune lineage subtype [14]. Low complexity cell transcriptomes with less than 90000 transcript reads and less than 1700 detected genes were filtered out (Fig. S1A and B). We used log2-transformed Counts Per Million (log2(CPM+1)) to allow comparison of read abundance across libraries of different sizes, unless otherwise specified. In an analysis subset, tumor-per-tumor data standardization was achieved by centering and reducing the data on a gene-by-gene basis (Datafile S1). Normalization on the basis of the expression of a set of 17 HKG is detailed in Supplementary Information 1. Tissue transcriptomes corresponded to the TCGA RNA-seq dataset of 155 untreated GBM patients [5] (normalized counts, log2(TPM+0.5), Table S1). Grouping analyses were performed using the Hierarchical Clustering on Principal Components (HCPC) approach (Datafile S1). Results were visualized using Principal Component Analysis (PCA) or t-distributed Stochastic Neighbor Embedding (tSNE). HCPC was also used to identify genes whose mean expression in one cluster differs from their mean across all cells (i.e. variables driving cell grouping). Mann-Whitney (Wilcoxon Rank Sum) test with p-values adjusted for multiple testing (Benjamini-Hochberg, p-value < 0.01) was used for differential gene expression analyses between cell or tissue groups [59]. R scripts used for unsupervised grouping and associated analyses are provided in Datafile S1. Normalized Mutual Information (NMI) scores were calculated to determine the contribution of cells issued from distinct tumors to each cluster. A NMI value of 1 implies that clusters gather objects (here, cells) corresponding to a single label (here, the tumor label) whereas a value of 0 denotes that all labels are split across all clusters (Table S1, [44]). Tumorigenic scores were obtained by computing the geometric mean of expression values of the signature genes per cell. When null, expression values were imputed a value of 1. Each gene was detected in at least 25% of GBM cells. Gene ontology analysis was done using the human genome as background (Table S1). Functional gene network reconstruction was achieved using the information-theoretic method, MIIC (multivariate information-based inductive causation, Supplementary Information 1 [58, 65]). Four independent datasets comprising 153 to 485 primary GBM transcriptomes were used for patient survival analysis (Table S1). Construction of a principal curve was achieved with a PCA based on the expression of each component of the lipid subgroup (16 genes coding for enzymes of the lipid metabolism overexpressed in Tum^HIGH^ cells and tissues and detected in at least 25% of GBM cells, see results). Each cell was projected onto the curve using the Pathifier algorithm (Table S1, [17]). Lipid subgroup and vesicle scores (Table S1) were calculated as described for the tumorigenic score.

### Biological experiments

Patient-derived cells (PDC) 6240**, R633 and 5706** were obtained from neurosurgical biopsy samples of distinct primary GBM, characterized and cultured as described [18, 57]. Lentiviral transduction (Table S1) was achieved as described [4, 18]. Viable cell counting, gene expression analysis, intracranial xenografts, and bioluminescence imaging (Table S1) were performed as described [4, 18]). All experiments were performed using independent biological samples, each independently repeated at least three times with the exception of the xenograft experiments. Prism 7.0 software (GraphPad) was used for statistical analyses with significance level set at p < 0.05.

## Results

### Unsupervised clustering analysis highlights first GBM cells’ tumor of origin

We used the publicly available single cell transcriptome dataset from Darmanis and colleagues [14] after removing low-complexity cell transcriptomes. Genes detected in at least 3 transcriptomes were retained for analysis (18577 genes for GBM cells and 19699 genes for normal cells). Previous analyses of GBM and other cerebral tumors scRNA-seq focused on the most dispersed [14] or most expressed genes [22, 50, 62, 64]. These analyses resulted in the identification of cell lineages and cell genomic anomalies rather than cell functioning states. We chose therefore to conserve all potential information by analyzing the full set of selected genes.

Gene expressions were computed as log2(CPM+1). Hierarchical Clustering on Principal Components (HCPC) of gene expressions in the mixed pool of GBM and normal cells readily separated cancer from normal cells (Fig. S1C). This result obtained by analyzing all detected genes is similar to the one obtained previously by analyzing the top 500 overdispersed genes [14]. Separate HCPC of normal cells resulted in six cell groups of immune or neural subtypes (astrocytes, oligodendrocytes, oligodendrocyte precursor cells, neurons, myeloid cells, vascular cells, Fig. 1b_1_), each group mixing cells from different tumors (Fig. 1b_2_). In striking contrast, GBM cell clusters resulting from HCPC analysis were dominated by cells from a single patient tumor (Fig. 1c). Contribution of the different tumors to clusters was scored by computing Normalized Mutual Information (NMI) between clusters and tumor labels, NMI scores being expected null if each tumor contributes equally to each cluster (Supplementary Material). Fully supporting our observation, the NMI score of normal cell clustering was only of 0.12 whereas the one of tumor cells was of 0.55 (two first bars in Fig. 1d). This predominant grouping of cancer cells by their tumor of origin has been reported for a number of cerebral and non-cerebral tumors [11, 14, 33, 37, 50, 54, 61]. For identifying traits common to all tumors, data can be analyzed tumor per tumor [62, 63, 69], or merged and analyzed as a whole after standardization (i.e. subtracting from each expression value the gene expression mean and dividing by its standard deviation across cells within a given tumor) [48]. The numbers of cancer cells precluding confident per-tumor analysis, we turned to standardization. HCPC analysis of standardized data resulted in clusters mixing cells from different GBM (Fig. 1e), as shown by a NMI score similar to the one calculated for normal cell grouping (third bar in Fig. 1d). However, gene ontology analysis of the genes identified in the HCPC as driving the cell grouping (see methods) did not provide clear links between cell clusters and potential cell functioning states (Fig. 1f, Table S2).

These results prompted us to further explore non-standardized data for minimizing the factors that might account for predominant grouping of cancer cells according to their tumor of origin.

### Tumor identity encoded by information dispersed through GBM cell transcript repertoires

Two main factors can account for tumor-driven cell grouping: the technical variations in tumor sample scRNA-seq, collectively referred to as batch effect, and the biological tumor-specific variations.

In scRNA-seq experiments, batch variations in RNA quality and sequencing efficiency, regardless of their origin, translate into variations in sample-dependent gene detection failures (referred to as dropouts) and in sample sequencing depth. Grouping of normal cells independently from their tumor of origin indicates that such batch variations are minor. We tested the influence of dropouts and of an additional normalization of the sequencing depth using a set of GBM-specific housekeeping genes (HKG) on the cell grouping (Table S3). Neither dropout imputation nor HGK normalization corrected tumor-driven cell grouping (Fig. 1d, Fig. S2A and B), confirming that batch effects are not major contributors of this grouping. Inter-tumor biological differences encompass genomic alterations known to vary greatly from one GBM to another, the tumor developmental stage, or the brain area and/or cells from which it developed [55]. We reasoned that differing biological characteristics, whatever their source, would translate into gene repertoires differing between tumors. To test this possibility, we considered binarization of the data by applying a value of one to all expressed genes regardless of their relative expression levels, and zero for non-detected ones. Maintenance of cell grouping by tumor of origin following binarization of gene expression (Fig. 1d and Fig. S2C) showed that cell gene repertoires are more similar within a given tumor than between two different tumors. We therefore sought to better understand which genes contribute most to this variability between cell transcriptomic landscapes. We first tested the impact of chromosome CNV on the cell grouping by filtering out genes mapped to chromosomes with CNV as previously identified [14]. Taking into account CNV did not modify tumor-driven cell grouping (Fig. 1d and Fig. S2D). Likewise, excluding genes detected in a single tumor or including only genes detected in all tumors did not change the outcome of HCPC analyses (Fig. 1d and Fig. S2E and F). We then tested the influence of inter-tumor variability in gene expression. Exclusion of the 100 genes identified as differentially expressed between tumors by Darmanis and colleagues [14] did not modify the outcome of the analysis (Fig. 1d and Fig. S2G). We then considered the most expressed genes, calculating the aggregate expression of each gene across cells, and retaining genes with the highest aggregate expression as described [62]. HCPC using the resulting 9505 most expressed genes still grouped cells by their tumor of origin (Fig. 1d and Fig. S2H). Likewise, performing HCPC using the top 500 or 1000 genes with expression variability between cells higher than expected (i.e. overdispersed [20]), or after excluding them, resulted in a similar tumor-driven cell grouping (Fig. 1d and Fig. S2I-L). Altogether, these results indicate that tumor-driven cell grouping is not based on limited and tumor-specific sets of genes. This led us to envisage that primary grouping of cancer cells by tumor of origin could result from information dispersed throughout the whole cell transcriptome. We challenged this hypothesis by performing iterative HCPC analyses on decreasing numbers of genes randomly selected among the 18577 detected genes. Ten analyses of distinct sets of randomly selected genes were performed for each size of gene sets (n = 2000, 1000, 500, 250, 100, and 50). NMI scores of the clusterings remained unchanged for gene set sizes > 500 (Fig. 2a and b). Their gradual decrease below this threshold indicated a progressive reduction of the influence of the tumor of origin on cell grouping. This influence was suppressed only when reducing the number of analyzed genes to 50, as shown by NMI scores equivalent to that calculated from the grouping analysis of the 19699 genes detected in normal cells (Fig. 2a and c). Altogether, these results show that tumor-driven cell grouping is irreducible to differential expression of circumscribed gene groups. To the contrary, it is encoded by information dispersed throughout the cell transcript repertoires, which is retrieved as soon as a combination of expressions of more than 500 genes is included in the analyses. As a consequence, unsupervised analysis turns out to be inadequate for identifying cell functioning states common to all tumors. We thus turned towards an approach of data reduction based on a signature of a functionally coherent set of genes.

**Fig. 2.**
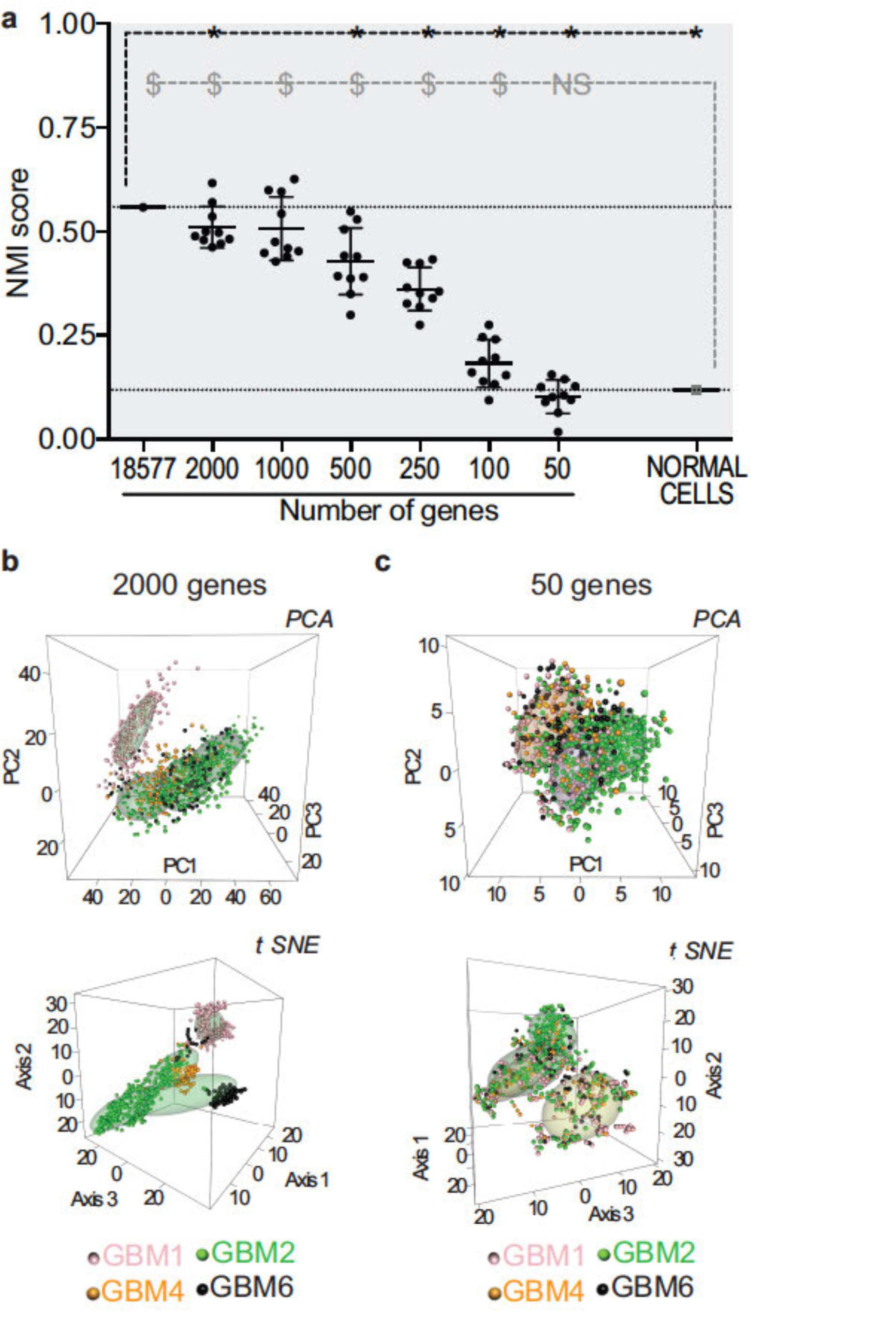
Down-sampling gene numbers relieves tumor-driven cell grouping. **a.** Decreased Normalized Mutual information (NMI) score when reducing gene numbers used for grouping analyses. Ten independent analyses performed with randomly selected genes for each gene number analyzed. * p <0.01 compared to the NMI score of the grouping analysis performed with all genes detected in GBM cells. $ p<0.0001 compared to the NMI score of the grouping analysis performed with all 19699 genes detected in the dataset of normal cells. One-sample t-test. Note that NMI scores consistently decrease below 500 genes analyzed, reaching values similar to the NMI score of the grouping analysis of normal cells only in grouping analyses performed with 50 genes. **b.** Example of a cell grouping analysis using 2000 randomly sampled genes. The clusters are predominantly composed of cells from a single tumor. **c.** Example of cell grouping analysis using 50 randomly sampled genes. Cells from a given tumor are distributed in different clusters. Each dot represents a cell colored according to its tumor of origin. Cell clusters delineated by ellipses.

### GBM cell grouping according to their tumorigenic potential upon signature-driven reduction of scRNA-seq data

We developed a grouping method based on a molecular signature we previously identified [4]. The signature is composed of five transcription factors, ARNT2, POU3F2, OLIG2, SOX9 and SALL2, with co-varied expression in GBM tissues and cells [4]. The POU3F2, OLIG2 and SOX9 genes have regulatory elements with ARNT2 binding sites and are down-regulated upon ARNT2 knockdown [4]. Each signature element was demonstrated to be required for GBM cell tumorigenic properties [4, 28, 41, 60]. We sought to use this signature to highlight subpopulations of cells in a tumorigenic state, expected to be present in all tumors.

To obtain an index of the cells’ tumorigenicity, we calculated a tumorigenic score corresponding to the geometric mean of the expression of each signature element. The score distribution curve exhibited a main inflection point corresponding to the distribution’s mean (Fig. 3a), which delineated two groups of 654 and 379 cells with low and high tumorigenic scores, respectively, hereafter designed as Tum^LOW^ and Tum^HIGH^ (Fig. 3a). Each of the four tumors contributed to each group (Fig. 3b, NMI score=0.057). Tum^HIGH^ GBM cells exhibited higher numbers of transcripts and genes than Tum^LOW^ GBM cells (Fig. S3A and B). We identified 6630 genes differentially expressed between both groups (Mann Whitney, BH-adjusted p-value < 0.01, Table S4), 98% of these genes being overexpressed in Tum^HIGH^ GBM cells. Of note, several items of the list of genes with enhanced expression in Tum^HIGH^ cells encoded proteins previously implicated in GBM cell aggressiveness (e.g. E2F1 [67], EGFR [1], HES1 and NOTCH1 [7], FABP7 [15], PTPRZ1 [24]). Conversely, genes with known tumor-suppressor properties were identified among the genes overexpressed in Tum^LOW^ cells (e.g. TUSC3 [34], SERPINB1 [31]). The whole workflow is summarized in Fig. S4 and provided in Datafile S2. These results suggest that cell functioning state can be inferred from scRNA-seq data following signature-driven data reduction. We next challenged the relevance of this approach by applying it to an independent dataset and using in vitro and in vivo GBM models.

**Fig. 3.**
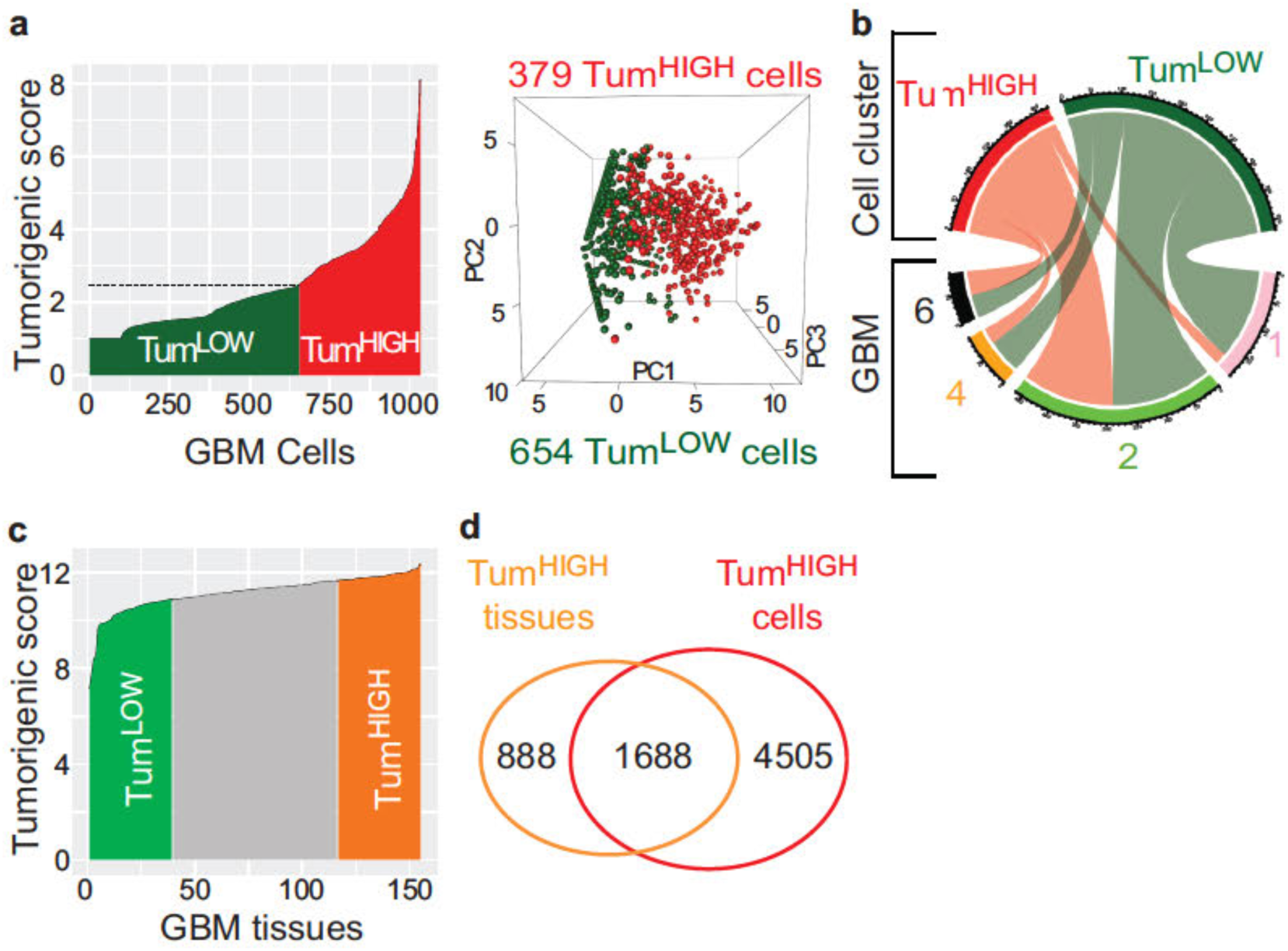
Signature-driven data reduction approach identifies cells according to their potential tumorigenic state. **a.** Splitting cells into groups with high (Tum^HIGH^) or low (Tum^LOW^) tumorigenic potential. Left panel: tumorigenic score distribution across the cells. Dotted line: mean of the tumorigenic score. Right panel: PCA plot based on the expression of the 5 elements of the tumorigenic signature. **b.** Contribution of each tumor to the two tumorigenic groups identified (chord diagram). Note that each tumor contributes to each cell group. **c.** Tumorigenic score distribution across GBM tissues (155 GBM tissues, TCGA RNA-seq dataset). Tum^HIGH^ and Tum^LOW^ GBM tissue groups selected at the extreme quartiles of the distribution. **d.** High overlap between genes overexpressed in Tum^HIGH^ GBM tissues and cells. 65.5% (1688) of genes overexpressed in Tum^HIGH^ GBM tissues are also overexpressed in Tum^HIGH^ GBM cells.

### Specific amino acid and lipid metabolic pathways distinguish GBM cells and tissues with high tumorigenic potential

To probe the biological relevance of our approach, we first applied the same analytical strategy to an independent dataset. Additional GBM single cell transcriptome datasets equivalent to the one published by Darmanis and colleagues and including several patients are not publicly available. We therefore turned to the TCGA collection of transcriptomes obtained by sequencing the RNA extracted from 155 patients’ GBM tissue fragments. As expected with respect to the heterogeneous nature of GBM tumors where cancer cells with differing properties co-exist with normal neural, vascular and immune cells, we observed a smoother distribution curve of the tumorigenic score across GBM tissues than across GBM cells (Fig. 3c). We therefore used quartiles to delineate two GBM tissue groups with low and high tumorigenic scores, respectively (Fig. 3c). Differential expression analysis between these two groups yielded a list of 6565 genes, 44% of them being overexpressed in Tum^HIGH^ GBM tissues (Mann Whitney, BH-adjusted p-value < 0.01, Table S4). The list of genes overexpressed in Tum^HIGH^ GBM tissues showed a 65.5% overlap with the list of genes overexpressed in Tum^HIGH^ cells (Fig. 3d, Table S4). This result was remarkable considering that it was obtained by confronting a dataset derived from 4 tumors to another derived from 155 tumors, and that tissue transcriptomes correspond to gene expression levels averaged over several hundred thousands of cells. Of the 1688 genes overexpressed in Tum^HIGH^ GBM cells and tissues, 78 encoded metabolic enzymes (Table S5). We further selected the 66 of them significantly correlated to the tumorigenic score across all GBM cells as well as across all GBM tissues (Pearson correlation, p-value <0.01, Table S5). Gene ontology analysis highlighted a 15 to 45-fold enrichment first in components of the lipid metabolism (27 genes) and second in components of the amino acids metabolic pathways (18 genes) (Fig. 4a, Table S5 and Fig. S5). Seven of the components of the amino acid metabolism belonged to the glycine, serine and threonine metabolism (Fig. 4b) whereas the lipid metabolism components were distributed among eight subpathways (Fig. 4b). Modeling of the regulatory gene network from the single cell expression data of the 66 metabolism genes using the MIIC algorithm singled out very long chain polyunsaturated fatty acid (VLC-PUFA) synthesis by highlighting ELOVL2 as the densest node of the network (Fig. 4c).

**Fig. 4.**
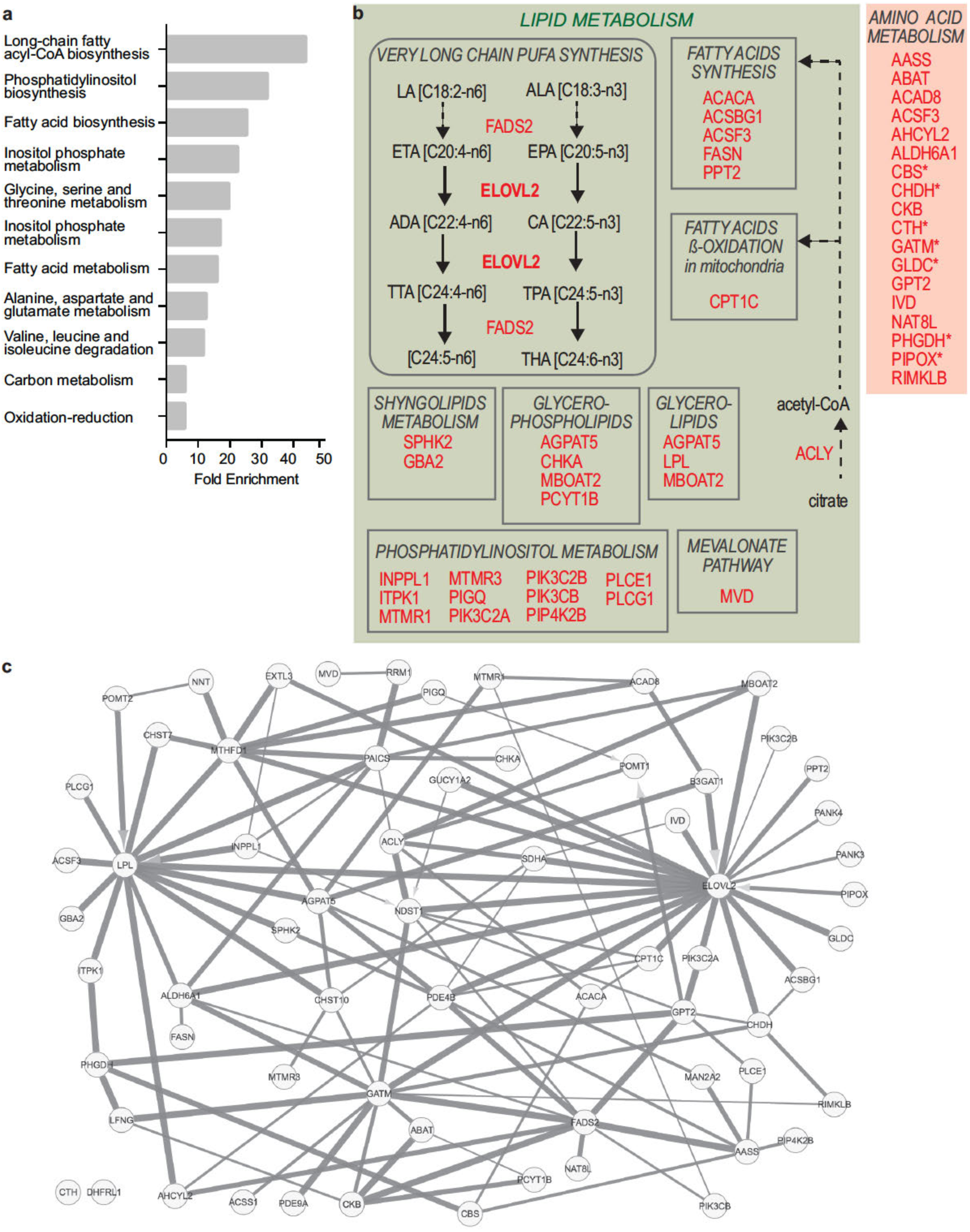
Enriched expression of genes coding for enzymes of the lipid and amino acid metabolisms in Tum^HIGH^ GBM cells and tissues. **a.** Gene ontology analysis of the 66 metabolism genes identified among genes overexpressed in both GBM Tum^HIGH^ cells and tissues. DAVID toolkit. **b.** Schematic representation of the lipid and amino acid metabolic pathways containing genes overexpressed in Tum^HIGH^ GBM cells and tissues. Asterisks mark genes coding for components of the glycine, serine and threonine metabolism. LA: linoleic acid; ALA: linolenic acid; ETA: eicosatetraenoic acid; EPA; eicosapentaenoic acid; ADA: docosatetraenoic acid; CA: clupanodonic acid; TTA: tetracostatetraenoic acid; TPA: tetracosapentaenoic acid; THA: tetracosahexaenoic acid. **c.** Modeling interconnections between the 66 metabolism genes highlights ELOVL2 at the most densely connected node of the network. Gene network built on the basis of the gene expression values across all GBM cells using MIIC tool. Line thickness represents the strength of the edge. Arrowheads linking variables in a v-structure of the type x → y ← z denotes the absence of a path between the x and z variables going through the y variable located on the tip of the v-structure.

### Functional association of the lipid metabolism enzyme ELOVL2 to patient clinical outcome and GBM development

ELOVL2 is a fatty acid elongase involved in the elongation of 22- to 24-carbon VLC-PUFA [27]. In agreement with ELOVL2 overexpression in Tum^HIGH^ cells, we observed ELOVL2 overexpression in cells with tumorigenic properties in an independent transcriptome dataset of human GBM cells in culture [38] (Fig. Fig. S6A). In addition, we observed a higher expression of ELOVL2 in GBM than in normal brain tissues (Fig. S6B), as well as in single GBM cells compared to single normal cells (Fig. S6C). Its expression was also higher in primary GBM characterized by a wild-type form of IDH1 compared to diffuse glioma, which are characterized by a mutant form of IDH1 (Fig. S6D). Of note, ELOVL2 levels were also higher in GBM bearing an amplified EGFR gene than in GBM with non-amplified EGFR (Fig. S6E). Finally, ELOVL2 high expression was found to be associated with worse patient survival in independent patient cohorts (Fig. 5a and Fig. S6F). Knocking down ELOVL2 expression in patient-derived cells (PDC) using lentiviral transduction of small hairpin (sh) RNA (Fig. 5b) resulted in a sharp decrease in cell proliferation (Fig. 5c). ELOVL2 role in the control of GBM cell tumorigenicity was evaluated in vivo using orthotopic xenografts of PDC stably expressing luciferase and either shControl or shELOVL2. Tumor development monitoring with bioluminescent imaging showed delayed tumor formation and reduced tumor burden in mice grafted with shELOVL2-PDC, compared to mice grafted with shControl-PDC (Fig. 5d and e). Of note, tumors that developed in a delayed manner from xenografts of shELOVL2-PDC had escaped from ELOVL2 inhibition, as shown by Q-PCR detection of human ELOVL2 mRNA levels at levels similar to those measured in tumors developing from xenografts of shControl-PDC (Fig. 5f). To gain insight into the cell process affected by ELOVL2, we selected from the 27 genes of the lipid metabolism, a lipid subgroup of 16 genes detected in at least 25% of the cells (Table S5). These genes were used to construct a principal curve onto which each GBM cell was projected (Fig. 5g). In addition, we computed a score with the expression of these 16 genes, following the same procedure as for the tumorigenic score. Of note, highest lipid scores (Fig. 5h), highest ELOVL2 expression values (Fig. 5i) and highest tumorigenic scores (Fig. 5j) all coincided in cells along the principal curve. This result further strengthens the relationship between the expression of the lipid subgroup, ELOVL2 expression and the tumorigenic state of the cells. Another member of the ELOVL family, ELOVL4, was involved in extracellular vesicle formation and release [29]. This prompted us to determine whether ELOVL2 expression is associated with molecular signatures of extracellular vesicles at the single cell level. Scores calculated for each of the four molecular signatures associated with extracellular vesicle were correlated with the tumorigenic scores (Table S6) and coincided with high ELOVL2 expression in cells along the principal curve (Fig. 5k and Fig. S7). This experimental set of results demonstrates that ELOVL2 is required for the tumorigenic behavior of GBM cells. It also suggests that ELOVL2 requirement stems from its involvement in the regulation of intercellular communication via extracellular vesicles. In addition, it provides robust experimental support for the relevance of signature-driven reduction of single cell transcriptomes to decipher the metabolic pathways underscoring GBM cell behaviors within the patients’ tumors.

**Fig. 5.**
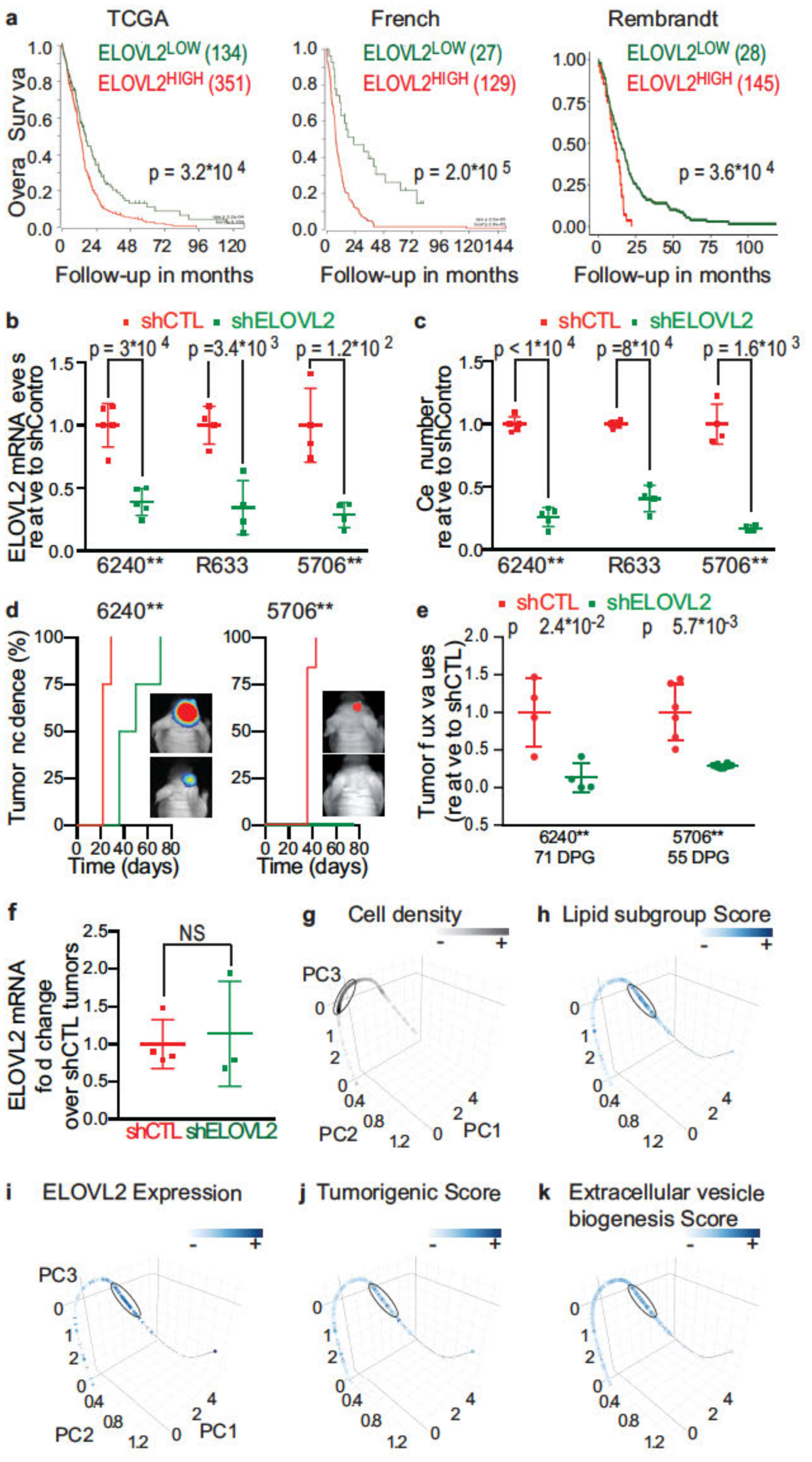
Association of the lipid metabolism enzyme ELOVL2 to patient clinical outcome and GBM development. **a.** High ELOVL2 expression is associated with worse patient survival. GBM tissue transcriptomes (microarrays) of 485, 156 and 173 GBM of the TCGA, French and Rembrandt datasets, respectively. Log-rank (Mantel-Cox) test. **b.** Decreased ELOVL2 mRNA levels in shELOVL2 patient-derived cells (PDC) compared to shControl cells. 6240** (left panel), R633 (middle panel) and 5706** (right panel) PDC. QPCR assay. Unpaired t-test with Welch’s correction, mean ± SD, n = 4-5 independent biological samples. **c.** ELOVL2 down-regulation decreases cell proliferation. 6240** (left panel), R633 (middle panel) and 5706** (right panel) cells. Unpaired t-test with Welch’s correction, mean ± SD, n = 4-5 independent biological samples. **d.** Knocking down ELOVL2 delays tumor development. Bioluminescent analyses of tumor growth initiated by grafting PDC transduced with a luciferase construct and either shControl (shCTL) or shELOVL2 constructs. n = 4 (6240**) and n = 6-8 (5706**) mice per group. **e.** ELOVL2 knockdown decreases tumor burden as shown by quantification of the tumor bioluminescent signals. DPG: days post-graft. n = 4 (6240**) and n = 6-8 (5706**) mice per group. Mean ± SD. Unpaired t-test with Welch’s correction. **f.** Recovery of ELOVL2 expression in tumors forming from xenografts of 6240** shELOVL2. QPCR assay. Mean ± SD. n=4 for shCTL and n=3 for shELOVL2. One sample t-test. **g-k.** Principal curve resulting from PCA of the expression of the subgroup of genes encoding lipid metabolism enzymes overexpressed in Tum^HIGH^ cells and tissues. **g.** Cell density along the principal curve. The ellipse delineates the portion of the curve with the highest cell density. **h-k.** Cells colored according to their (h) score calculated with the components of the lipid subgroup, (i) ELOVL2 expression levels, (j) tumorigenic score, and (k) extracellular vesicle biogenesis score. Note that cells with either high score or expression value cluster on the same portion of the curve (ellipses). Pathifier tool.

## Discussion

Metabolism is at the heart of cell behavior and its dependence on enzymatic activities makes it a target of choice for therapeutic targeting. GBM cells adapt their functioning state to changes in their microenvironment, whether these changes result from natural tumor growth or therapies. Progress towards identification of the most relevant possible targets for patient therapy requires access to molecular players active in the actual context of the patient tumors while taking into account the heterogeneity of the tumor tissue. Transcriptomes of single cells sorted from patient GBM offer such an access.

Unsupervised analysis of scRNA-seq resulting in predominant grouping of cancer cells according to their tumor of origin and of normal cells independently from their tumor of origin was found in analyzing scRNA-seq from tumors other than GBM [11, 33, 37, 54, 61] but never questioned. We considered in all ways we thought eventual influences of scRNA-seq technical biases as well as of the tumor-specific differences in gene repertoires expected to reflect inter-tumor biological differences whatever their sources. We did not find one accounting for cell grouping-dependence on the tumor of origin. In contrast, unsupervised analyses of randomly down-sampled numbers of genes showed that tumor-driven cell grouping disappears below a critical number of genes (100 to 50 in this study). Collectively, our results show that cancer cell clustering per tumor is not due to scRNA-seq technical bias (as expected with respect to the lack of influence of the tumor on normal cell grouping) or to circumscribed gene subsets reflecting inter-tumor biological differences. They support the notion that the identity of the tumor to which each cancer cell belongs is encoded by information dispersed throughout the cell transcript repertoire.

To unmask cell functioning states regardless of their tumor of origin, we reduced the data based on previously acquired biological knowledge. Signature-driven data reduction was previously used to infer cell lineages from single cell transcriptomes of oligodendrogliomas and diffuse infiltrative pontine gliomas [22, 62]. In these studies performed on centered data, cell lineages were inferred using sets of top correlated genes to the principal component scores of a PCA of the dataset under scrutiny [62], or on the basis of mouse or human gene sets differentiating normal neural subtypes [22, 64]. Here, we based our analysis on a molecular signature related to a major functioning state, tumorigenicity.

Cells from a given GBM have long been known to be endowed with differing abilities to initiate neoplasms [30, 53, 66]. We postulated that such a choice would enhance the likelihood to group cells based on their functioning state irrespective of their tumor of origin. We postulated also that using a signature defined [4] and experimentally validated in independent prior studies [4, 28, 41, 60] would reduce the risk of obtaining results relevant only for the dataset analyzed. Our choices proved fruitful to identify two contrasting functioning states, with respect to the following findings: genes previously described as controlling GBM cell aggressiveness were found to be overexpressed in cells with high tumorigenic scores; a highly similar list of differentially expressed genes was obtained when applying the same analytical strategy to an independent dataset made of tissue bulk transcriptomes of a much larger number of GBM; highest scores for the lipid subgroup found to be enriched in Tum^HIGH^ cells and tissues coincided with highest tumorigenic scores; finally, experimental knockdown of the most interconnected gene of this lipid metabolism subset, ELOVL2, impaired GBM cell tumorigenic properties.

Oncogenic mutations have long been known to favor mobilization of metabolic pathways, most notably by allowing cancer cells to adapt their mode of energy production to different microenvironments and sources of nutrients [40, 52]. GBM cells, like other cancer cell types, are considered to favor glycolysis over oxidative phosphorylation for ATP production. This prevalence of glycolysis is considered as a cell adaptation to its need for feeding carbon into biosynthesis of nucleic acids, fatty acids and proteins for cell growth and proliferation. At the single cell level, overexpression of genes coding for enzymes of the glycolysis pathway was not associated with high tumorigenic scores. This suggests that enhanced mobilization of glycolysis enzymes is not proper to GBM cells in a tumorigenic state, at least at the transcriptional level. Overexpression of two genes coding for enzymes of the amino-acid pathways, GATM and CKB (Fig. 4b) rather suggests that another source of energy distinguishes tumorigenic from non-tumorigenic GBM cells. GATM is one of the two enzymes ensuring brain-endogenous synthesis of creatine, starting from glycine. The second enzyme, GAMT, was found to be overexpressed in tumorigenic high GBM cells but not retained for final analyses because absent from the list of genes overexpressed in tumorigenic GBM tissues. CKB is responsible for the phosphorylation of creatine, which serves for ATP regeneration and plays an essential role in brain energy metabolism [56]. Interestingly, GATM is the most interconnected of the enzymes coding for elements of amino acid metabolism in the modeling of the gene expression network (Fig. 4c). Lipids are also a significant source of energy. Lipid metabolism association to GBM cell aggressiveness has been reported to stem from lipid contribution to cell energetics through fatty acid beta-oxidation and to transduction pathways through the mevalonate metabolism [40]. Accordingly, we found that genes coding for key enzymes of the fatty acid synthesis (e.g. ACLY, FASN) and beta-oxidation pathways (CPT1), as well as the mevalonate pathways (MVD) were overexpressed in Tum^HIGH^ cells. In addition, we observed an overexpression of genes involved in the synthesis of phospholipids, glycerolipids and sphingolipids, essential components of plasma membranes and/or sources of potent autocrine/paracrine signaling molecules.

The biological relevance of the results of our bioinformatics analyses was further validated in in vivo experimental models of human GBM. Our study unveiled an unexpected causal link between ELOVL2, the endpoint enzymatic component of the lipid subpathway ensuring synthesis of VLC-PUFA, and the tumorigenic status of GBM cells. Notably, we show that ELOVL2 knockdown in PDC decreases tumor growth in vivo. Little is known on this member of the ELOVL family that catalyzes the elongation of saturated and monounsaturated VLCFA (ELOVL1, 3, 6 and 7) and of VLC-PUFA (ELOVL2, 4 and 5) by adding two carbon units to the carboxyl end of a fatty acid chain [27]. ELOVL2 is specifically involved in the elongation cascade starting from the dietary PUFA linoleic and linolenic acids, which cannot be synthesized by humans. The products of the enzymatic process of these essential PUFA are thought to modulate diverse biological phenomena ranging from cell survival to inflammatory responses [3, 10]. In mouse models, Elovl2 knockout has been shown to result in defective PUFA composition in the liver, serum and testis in association with male infertility and a reduced capacity to accumulate fat [51, 70]. PUFA are structural components of membrane phospholipids especially enriched in neural tissues, and provide potent signaling compounds [39]. In the MCF7 cell line that models breast cancers requiring estrogen for growth, ELOVL2 expression is positively regulated by estrogen [26], and its knockdown is associated with epithelial to mesenchymal transition [36]. In this epithelial cancer high ELOVL2 expression is associated with higher metastatic relapse-free survival [36]. On the opposite, ELOVL2 upregulation in prostate cancer has been associated with the oncogenic effect of SPOP loss of function mutations [68]. Here, finding of the association between high ELOVL2 expression in GBM and worsened patient prognosis (Fig. 5a and Fig. S6F), coupled with the demonstrated requirement of ELOVL2 for GBM cells tumorigenicity in vivo, demonstrates a causal link between ELOVL2 and GBM growth. We investigated through bioinformatics analysis what might be the mechanism of action of ELOVL2 overexpression in tumorigenic high cells. The correlation between ELOVL2 overexpression and molecular signatures of extracellular vesicles suggests that formation and release of extracellular vesicles is one of the cell processes by which ELOVL2 controls GBM tumor development. This possibility is coherent with the reported involvement of another member of the family, ELOVL4, in the formation of synaptic vesicles in the brain and retina [29]. Extracellular vesicles have been involved in intercellular communications within GBM, by carrying metabolites, nucleotides and proteins able to affect the behavior of cancerous as well as non-cancerous cells composing the tumor [6, 21]. Our experimental results are coherent with our modeling results that place ELOVL2 at the core of the metabolic pathways essential for sustaining GBM cell tumorigenicity. ELOVL2 importance for GBM is strengthened by a study published during writing of this article that describes ELOVL2 as a super-enhancer associated gene controlling glioblastoma stem cell properties [25].

## Conclusions

The present findings underscore the power of single cell transcriptome analyses for unveiling the complexity of the participation of metabolism in relation to the heterogeneity of cell functioning states encountered in GBM. It is worth emphasizing that the discovery of a molecular deregulation that proved to be a predictor of patient survival in independent cohorts of several hundred tumors stems from the study of cells derived from only four tumors. Our results show the high relevance of integrating the cell functioning status, even when focusing on only two contrasting states, for the discovery of metabolic modules controlling GBM aggressiveness. Further development of signature-driven data reduction based on established experimental evidence will lead to further refine the identification of functioning states and of the diversity of the molecular networks required for their maintenance. The workflow we designed to classify any single cell dataset according to a signature score will be instrumental for blocking GBM cell ability to escape therapies, thus contributing to improve GBM therapies.

## List of abbreviations

CNV: Copy Number Variation
CPM: Counts Per Million
GBM: Glioblastoma
HCPC: Hierarchical Clustering on Principal Components
HKG: HouseKeeping Genes
MIIC: Multivariate Information-based Inductive Causation
NMI: Normalized Mutual Information
PCA: Principal Component Analysis
PDC: Patient-Derived Cells
scRNA-seq: single cell RNA-sequencing
shRNA: small hairpin RNA
tSNE: t-distributed Stochastic Neighbor Embedding
Tum: Tumorigenic
VLC-PUFA: Very Long Chain PolyUnsaturated Fatty Acids

## Declarations

### Ethics approval and consent to participate

All the procedures performed in studies involving human participants were in accordance with the 1964 Helsinki declaration and its later amendments and to the French laws. The institutional review board of the Sainte-Anne Hospital Center - University Paris Descartes (Comité de protection des personnes Ile de France III) approved the study protocol (Protocol number DC-2008-323). All samples were obtained with informed consent of patients. All animal maintenance, handling, surveillance, and experimentation were performed in accordance with and approval from the Comité d’éthique en expérimentation animale Charles Darwin N°5 (Protocol #5379).

### Consent for publication

Not applicable.

### Availability of data and material

All data are provided in the manuscript.

### Author contributions

Conceptualization, GMD, HC and MPJ; Methodology, MSS, LB, GMD, CA, HC and MPJ; Experimental investigations, MSS, EAE, VD; Computational analyses, MSS, LB, CA, GMD; Writing – Original Draft, MSS and MPJ; Writing – Review & Editing, MSS, EAE, HC, CA, GMD, MPJ; Supervision, GMD and MPJ; Project Administration, HC; Funding Acquisition, HC, MPJ, CA.

## Acknowledgments

We are grateful to Nadir Sella (Institut Curie, CNRS UMR168, Paris) for his guidance and help for using MIIC, and to François-Xavier Lejeune (ICM, Salpétrière Hospital, Paris) for his advices on statistical analyses.

## Supplementary Information, tables and figures legends

**Supplementary Information 1** (related to Material and Methods).

**Datafile S1. R scripts used for unsupervised grouping and associated analyses.**

**Datafile S2. R scripts used for signature-based analytical workflow.** Schematic representation of the workflow in Fig. S4.

**Table S1** (related to Material and Methods). **List of all resources and materials, R packages, corresponding websites and references.**

**Table S2** (related to Fig. 1)**. Gene ontology (GO) analysis of the genes describing each of the seven clusters identified upon grouping analysis using standardized data.** DAVID toolkit. Sheet 1: Legend. Sheet 2: Lists of genes upregulated in a given group, as compared to their mean across all cell groups, and used for GO analysis. Following sheets: GO analysis results for clusters 1-7.

**Table S3. List of housekeeping genes used for data normalization.** Sheet 1: Legend. Sheets 2-3: Percentage of cells in which genes are detected (sheet 2), the coefficient of variation of their expression (sheet 2) and their auto-correlation (sheet 3) across all GBM cells are indicated. Computing gene expression values with the log2(CPM + 1) allows correcting sequencing depth variations between samples. We tested an additional step in the normalization of the sequencing depth based on a set of housekeeping genes (HKG). We considered as HKG the cluster of genes detected in at least 90% of the cells with pairwise correlated expression and the lowest coefficient of variation across cells. The geometric means of HKG expression were calculated per small groups of cells with similar expression profiles (metacells). Gene expression values computed as log2(CPM + 1) were divided, for each cell, by the normalization factor corresponding to its metacell.

**Table S4** (related to Fig. 3)**. Genes differentially expressed between low and high tumorigenic GBM cells and/or tissues. Sheet 1: Legend. Sheet 2:** Genes differentially expressed between GBM cells with low and high tumorigenic scores. **Sheet 3:** Genes differentially expressed between GBM tissues with low and high tumorigenic scores. **Sheet 4**: Genes found to be overexpressed in Tum^HIGH^ cells as well as tissues.

**Table S5** (related to Fig. 4)**. Pearson correlation across all GBM cells (sheet 2), and all GBM tissues (sheet 3) between the tumorigenic score, and the 78 metabolism genes identified as overexpressed in Tum**^**HIGH**^ **cells and tissues.** Sheet 1: legend. Sheet 4: List of the 66 genes significantly correlated to the tumorigenic score across all GBM cells as well as across all GBM tissues. Sheet 5: Gene ontology analysis of the 66 genes coding for metabolic enzymes. Sheet 6: List of the 27 genes coding for enzymes of the lipid metabolism. Genes detected in at least 25% of the cells are highlighted.

**Table S6** (related to Fig. 5 and Fig. S7)**. Correlation values calculated across all cells between the tumorigenic score, ELOVL2 expression, and the signature scores associated with extracellular vesicle biogenesis, transport, targeting, and fusion.**

**Fig. S1** (related to Fig. 1). **Filtering out cells with low-complexity transcriptomes and unsupervised grouping analysis of GBM and normal cells simultaneously.** A-B. Filtering out cells with low-complexity transcriptomes. Cells with more than 90000 transcripts and more than 1700 genes were selected for further analyses. 1033 GBM (A) and 2417 normal cells (B) were retained. Left panels: selected cells colored in blue, rejected cells colored in gray. Right panels: cells colored by tumor. C. Unsupervised grouping analysis of the mixed set of GBM and normal cells distinguishes cancer cells from normal cells. Each dot represents a cell. Normal cells colored in light blue, cancer cells colored in violet. Left panel: PCA visualization. Right panel: tSNE visualization.

**Fig. S2** (related to Fig. 1). **Maintenance of tumor-driven cell grouping regardless of the mode of data normalization or filtering.** A: Multidimensional scaling (MDS) visualization. B-L: PCA visualization. Each dot represents a cell. Ellipses delineate the clusters identified. **A-B.** Technical biases related to scRNA-seq do not contribute to tumor-driven cell grouping. **A.** Maintenance of tumor-driven cell grouping after imputation of dropouts using the algorithm CIDR (Clustering through Imputation and Dimensionality Reduction). CIDR imputes dropouts by inferring their values from gene expression across all cells [42]. **B.** Normalizing data by the expression of housekeeping genes (HKG) prior to analysis fails to alleviate tumor-driven cell grouping. **C-L.** Tumor-specific biological differences are not reducible to circumscribed sets of genes. **C.** Binarization of expression data to overcome possible inter-experimental variations in the efficacy of scRNA-seq, and therefore in the detected RNA levels, does not change tumor-driven cell grouping. **D.** Filtering out genes located on chromosomes with identified variations in copy number (CNV) does not modify tumor-driven cell grouping. **E.** Tumor-driven cell grouping is maintained after removal of genes detected in a single tumor. **F.** Grouping analysis performed using only genes detected in all tumors also results in tumor-driven cell grouping. **G.** Filtering out genes differentially expressed between tumors does not alleviate tumor-driven cell grouping. **H.** Cell grouping analyses using the most expressed genes across cells fails to overcome tumor-driven cell grouping. **I**. Cell grouping analyses using the top 500 genes with the most overdispersed expression across cells fails to overcome tumor-driven cell grouping. **J.** Filtering out the top 500 genes with the most overdispersed expression across cells does not change tumor-driven cell grouping. **K.** Cell grouping analyses using the top 1000 genes with the most overdispersed expression across cells fails to overcome tumor-driven cell grouping. **L.** Filtering out the top 1000 genes with the most overdispersed expression across cells does not change tumor-driven cell grouping.

**Fig. S3** (related to Fig. 3). Increased numbers of transcript (A) and genes (B) detected per cell with high tumorigenic scores, compared to cells with low tumorigenic scores. Mann-Whitney test. p = 1.92*10^−3^ for transcript number and 2*10^−41^ for gene number.

**Fig. S4. Signature-based analytical workflow.**

The analytical method developed has been implemented in R and is provided in Datafile S2.

**Fig. S5** (related to Fig. 4)**. Schematic representation of the main metabolic pathways in which are involved 60 of the 66 metabolic enzyme genes overexpressed in Tum**^**HIGH**^ **GBM cells and tissues.** Asterisks mark genes coding for components of the glycine, serine and threonine metabolism.

**Fig. S6** (related to Fig. 5)**. Increased ELOVL2 expression is associated with increased tumor burden. A.** Down-regulated ELOVL2 expression in patient-derived GBM cells deprived of tumorigenic properties, compared to their tumorigenic counterparts. Mann-Whitney test. Lee microarray dataset GEO ID: GSE4536. R2 genomics analysis and visualization platform database (http://r2.amc.nl). **B.** Higher ELOVL2 expression in GBM tissues compared to normal brain tissues. Mann-Whitney test. TCGA tissue transcriptome dataset (microarrays) of 528 primary GBM and 10 normal brain tissues. **C.** Significantly higher ELOVL2 expression in GBM cells compared to normal cells. Mann-Whitney test. scRNA-seq dataset from Darmanis and colleagues [14]. **D-E.** ELOVL2 expression prevails in GBM bearing a wild-type form of *IDH1* (D) and with *EGFR* gene amplification (*EGFR*^AMP^) (E). French tissue transcriptome dataset (microarrays). IDH1^WT^, n=95. IDH1^MUT^ n=33. *EGFR*^WT^, n=46. *EGFR*^AMP^, n=32. Mann-Whitney test. **F.** High ELOVL2 expression is associated with a poorer survival for patients. TCGA GBM tissue transcriptomes (RNA-seq) of 153 primary GBM. Log rank test.

**Fig. S7** (related to Fig. 5)**. Principal curve analysis associates GBM cell tumorigenic state to mobilization of vesicle production and release.** Principal curve resulting from PCA of the expression of the subgroup of genes encoding lipid metabolism enzymes overexpressed in Tum^HIGH^ cells and tissues. Cells colored according to their score calculated with the components of molecular signatures associated with extracellular vesicle transport (A), targeting (B), and fusion (C). Note that cells with either high score cluster on the same portion of the curve (ellipses).

