## Supplementary Materials for "Capture at the single cell level of metabolic modules distinguishing aggressive and indolent glioblastoma cells"

*Public data acquisition*

The single cell transcriptomes of 1091 GBM cells and 2498 normal cells originating from four patients were downloaded from http://gbmseq.org/ [14]. This dataset comprises only good-quality cells, as determined by its authors on the basis of the expression of 35 housekeeping genes (HKG), and distinguishes cancer from normal cells on the basis of CNV profiling. Normal cells are further distinguished according to their neural or immune lineage subtype [14]. Normalized counts in $\log_2$(TPM+0.5) from TCGA RNA-seq data of 155 untreated GBM patients with available survival data were obtained from the Gliovis portal (http://gliovis.bioinfo.cnio.es/).

*Standardization or normalization of single cell transcriptome data*

For analyses of single cell transcriptomes, we used log2-transformed Counts Per Million ($\log_2$(CPM+1)), unless otherwise specified. CPM corresponds to the counts of gene mapped reads normalized by the total number of mapped reads per cell library divided by one million, thus allowing comparison of read abundance across libraries of different sizes. To avoid potential analytical bias due to scarcely detected genes, we filtered out genes detected in less than 3 cells prior to grouping analysis, keeping 18577 genes for GBM cells and 19699 genes for normal cells. When grouping analysis included both GBM and normal cells, we kept the 20237 genes detected in at least 3 GBM cells or 3 normal cells.

In a subset of analyses, tumor-per-tumor data standardization was achieved by centering and reducing the data on a gene-by-gene basis as described: $x_{Ti}$'= ($x_{Ti}$ – mean($x_T$)) / sd($x_T$)), where $x_{Ti}$ corresponds to a gene transcript level in a given cell from a given tumor T, and mean($x_T$) and sd($x_T$) to the arithmetic mean and standard deviation across GBM cells of tumor T, respectively [8, 43]. Note that grouping analysis using data standardized by tumor resulted in 10 clusters, including 3 with only 1 cell each. These 3 clusters were ignored in further analyses.

In another subset of analyses, data were normalized on the basis of the expression of a set of housekeeping genes (HKG). The 35 genes considered by Darmanis *et al* as HKG were not used because of their variable detection in 2.13 to 99.9% GBM cells and variable expressions (coefficients of variation (CV) ranging from 0.10 to 7.93). Here, we defined HKG according to the following criteria: genes detected in most cells, with relatively constant expression across all cells and whose expressions auto-correlate. Thus, calculation of the Pearson correlation (corr) between each pair of genes was followed by calculation of the distance matrix between each pair of genes based on their correlation matrix (1-corr). Groups of genes with correlated expressions were identified using a hierarchical clustering (Ward's method using ward.D2 algorithm, cutting dendrogram into clusters at h=1.2). Next, we selected the gene group with the smallest average CV of gene expression. From this group, we retained only genes detected in at least 90% cells, defining thus a set of 17 HKG expressed with a CV of their expression ranging from 0.19 to 0.42. To normalize data according to HKG, we first defined small groups of 3 to 65 cells with similar expression profiles, using the Pearson correlation between each pair of cells and the corresponding distance matrix (1-corr), followed by hierarchical clustering (Ward's method using ward.D2 algorithm, with h varying between 0.8 and 1.1 to obtain clusters with less than 65 cells). The arithmetic mean expression of each HKG was calculated for each of these small cell groups considered as metacells. The normalization factor corresponded to the geometric mean of all HKG per metacell (psych R package for geometric mean calculation). The data were normalized by dividing each gene expression value per cell by the normalization factor of its corresponding metacell.

*Grouping analyses*

Grouping analyses were performed using the Hierarchical Clustering on Principal Components (HCPC) approach (FactoMineR package). This approach combines three standard methods for multivariate data analyses performed stepwise: Principal Component Analysis (PCA), hierarchical clustering, and partitioning clustering by the k-means method (http://factominer.free.fr/more/HCPC_husson_josse.pdf). PCA was performed to reduce the dimensions to the first 10 principal components (PCs). Euclidean method was used to construct a cell-to-cell distance matrix. Hierarchical clustering was then performed on this distance matrix using the Ward's criterion (ward.D2 algorithm). The resulting partitioning of the cells was improved by a K-means clustering with 10 iterations. The cell grouping was visualized using PCA (FactoMineR package) or tSNE implemented using the tsne R package (perplexity = 50). Graphs were generated with the rgl and magick

R packages. In addition, HCPC allows identifying the variables that drive cell grouping for each cell cluster, i.e. genes whose mean expression in one cluster differs from its mean across all cells (pval < 0.05, t-test).

A Normalized Mutual Information (NMI) score was calculated to determine the contribution of cells issued from distinct tumors to each cluster (ClusterR package). This metric is used as an external clustering validation that compares clustering results to a known truth [44]. A NMI value of 1 implies that clusters gather objects (cells) corresponding to a single label (here, the tumor label), whereas a value of 0 denotes that all labels are split across all clusters. Graphs were generated with Prism 7.0 software (GraphPad).

*Differential gene expression analysis*

Differential gene expression analyses between differing cell or tissue groups were performed using the Mann-Whitney (Wilcoxon Rank Sum) test [59]. The p-values were adjusted for multiple testing using Benjamini-Hochberg (BH) approach. The level of significance was set at BH-adjusted p-value < 0.01. Fold change (FC) for gene i was calculated as follows: $FC_i=x_i-y_i$, where $x_i$ and $y_i$ are the log2 expression levels of gene i in conditions x and y, respectively. Only genes detected in at least 3% of GBM cells or tissues were considered for this analysis. Genes coding for metabolism enzymes were identified using the list from KEGG [35].

*Gene ontology analysis*

Functional enrichment analysis was carried out using the online database DAVID v6.8 (https://david.ncifcrf.gov/). The human genome was used as background (Homo Sapiens from DAVID). We considered p-value < 0.05 (Fisher's Exact test) as the cut-off criterion for significance. Graphs were generated with Prism 7.0 software (GraphPad).

*Functional gene network reconstruction*

Prior to the network reconstruction, expression data were binarized into an ON-OFF system. For each cell, detected genes were considered as ON and assigned a value of 1 regardless of their relative expression levels. Undetected genes were considered as OFF and retained their original value of 0.

The information-theoretic method, MIIC (multivariate information-based inductive causation), was used for gene network reconstruction and performed on the dedicated website (https://miic.curie.fr/) [[65]; [58]]. This method is based on the analysis of multivariate information, which extends the concept of mutual information beyond two variables and may imply cause-effect relationships between the underlying variables. On the resulting graph, the edge between 2 variables X and Y reflects their connection with an edge-specific confidence ratio <0.01. The lower the ratio, the higher the confidence.

*Cell cultures*

Patient-derived cells (PDC) 6240**, R633 and 5706** obtained from neurosurgical biopsy samples of distinct primary GBM were cultured in defined medium containing bFGF and EGF as described [18, 57]. The cells were transduced with lentiviral vectors encoding a control or an ELOVL2 shRNA construct (pLKO.1-HPGK-puro-U6-non mammalian shRNA control, and pLKO.1-puro-CMV-tGFP-U6-shELOVL2-64, Sigma, France). Non-transduced cells were eliminated following puromycin treatment (2 µg/mL). Lentivirus was produced by the Plateforme vecteurs viraux et transfert de gènes (Necker Federative structure of research, University Paris Descartes, France).

*Viable cell counting*

Trypan blue exclusion test was used to determine the number of viable cells (Trypan blue solution, ThermoFisher, 0.4% v/v, 3 min incubation at room temperature). Blue and white cells (dead and alive, respectively) were counted with the Countess automated cell counter (Thermo Fisher, France).

*Gene expression analysis*

Total RNA was prepared using the Nucleospin RNA kit (Macherey-Nagel) according to the manufacturer's instruction. Contaminating DNA was removed using an rDNAse solution supplied with the Nucleospin RNA kit during RNA isolation. cDNA was prepared using the QuantiTect Reverse Transcription Kit (Qiagen) according to manufacturer's instructions. QPCR assays were performed using the LightCycler480 (Roche, France) and the SYBR Green PCR Core Reagents kit (Bimake.com). The thermal cycling conditions comprised an initial denaturation step at 94 °C for 5 min, and 40 cycles at 94 °C for 30 s, 60 °C for 30 sec and 72 °C for 30 sec. Transcripts of the TBP gene encoding the TATA box-binding protein (a component of the DNA- binding protein complex TFIID) were quantified as an endogenous RNA control. Quantitative values were obtained from the cycle number (Cq value), according to the manufacturer's manuals. Sequences of primers used for QPCR are: ELOVL2 - forward primer: 5'-TCCACTTGGGAAGGAGGCTACA; ELOVL2 - reverse primer: 5'-

CTCCAAATCAGTAGAGTTCCTGG; TBP - forward primer: 5'-TGCACAGGAGCCAAGAGTGAA; TBP - reverse primer: 5'-CACATCACAGCTCCCCACCA.

*Intracranial xenografts*

The animal maintenance, handling, surveillance and experimentation were performed in accordance with and approval from the Comité d'éthique en expérimentation animale Charles Darwin N° 5 (Protocol #5379). 6240** and 5706** PDC transduced with lentiviruses encoding either a shControl or a shELOVL2, were used. 20,000 cells (6240**) or 100000 cells (5706** ) were injected stereotaxically into the striatum of anesthetized 8-week-old Nude mice (Envigo Laboratories, France), using the following coordinates: 0 mm posterior and 2.5 mm lateral to the bregma, and 3 mm deep with respect to the surface of the skull. Bioluminescence imaging was performed on a Photon Imager Biospace (Biospace Lab, France), after intra-peritoneal injection of 150 µL luciferin (20 mM, Thermo Fisher, France). Tumor formation was monitored by bioluminescence until all mice of the control group showed a signal. Bioluminescent signals were visualized with M3 Vision software (Biospacelab).

MS Saurty, L Bellenger, E A. El-Habr, V Delaunay, H Chneiweiss, C Antoniewski, G Morvan-Dubois, MP Junier
Signature-driven single cell transcriptomes analysis captures metabolic modules distinguishing aggressive and indolent glioblastoma cells

**Table S1 (related to Material and Methods). List of all resources and materials, R packages, corresponding websites and references.**

| REAGENT or RESOURCE | DESCRIPTION or APPLICATION | SOURCE | REFERENCES |
|---|---|---|---|
| **Data Acquisition** | | | |
| scRNA-seq from human GBM cells | Single ce l transcriptomes from 4 GBM patients, Count values | http://gbmseq.org/ | [14] |
| RNA-seq from human GBM tissues | The Cancer Genome Atlas (TCGA) tissue transcritpomes from 155 untreated GBM patients, log2(TPM+0.5) values | Gliovis portal (http //g iovis.bioinfo.cnio.es) | [5] |
| DNA microarray data from RNA extracted from early passage (P3) of cultured GBM ce ls isolated from 4 human GBM. | Transcriptomes of cell cultures from GBM cell cultures with and without tumorigenic properties | R2 genomics analysis and visualization platform database (http //r2.amc.nl) | [38] |
| **Algorithms or Softwares**<br>*See Datafiles S1 and S2 for Detailed scripts* | | | |
| Adobe Illustrator | Figure preparation | Adobe Systems | |
| R language version 3.5.0 | The R Project for Statistical Computing | https://cran.r-project.org/ | |
| GraphPad Prism 7.0 software | Statistical analyses, graphs | GraphPad | |
| psych R package | Geometric mean calculation | https://personality-project.org/r/psych | |
| FactoMineR R package | Hierarchical Clustering on Principal Components (HCPC) for unsupervised grouping analyses; ward.D2 algorithm used to calculate the distance matrix between ce ls | http://factominer.free.fr/ | Husson F, Josse J, Pagès J, 2010 |
| tsne R package | tSNE visualization, perplexity 50 | https://github.com/jdonaldson/rtsne/ | |
| rgl R package | Plots of PCA or tSNE resu ts | https://r-forge.r-project.org/projects/rgl/ | |
| magick R package | Plots of PCA or tSNE resu ts | https://github.com/ropensci/magick#readme | |
| ClusterR R package | Normalized Mutual Information (NMI) score calculation | https://github.com/mlampros/ClusterR | [44] |
| ggplot2 R package | Score distribution and violin plots | http://ggplot2.tidyverse.org/ | |
| circlize R package | Circus plot | https://github.com/jokergoo/circlize | |
| DAVID version 6.8 | Functional enrichment analysis | https://david.ncifcrf.gov/ | |
| MIIC tool | Multivariate Information-based Inductive Causation; Gene network reconstruction | https://miic.curie.fr/ | [58, 65] |
| Pathifier algorithm | Principal curve analysis | www.weizmann.ac.il/pathifier | [17] |
| M3 Vision software | Visualisation of bioluminescent signals | Biospace Lab, France | |
| **Data standardization approach** | Tumor-per-tumor data standardization was achieved by centering and reducing the data on a gene-by-gene basis as described xTi' (xTi – mean(xT)) / sd(xT)), where xTi corresponds to a gene transcript level in a given cell i from a given tumor T, and mean(xT) and sd(xT) to the arithmetic mean and standard deviation across GBM cells of tumor T respectively. | | [8, 43] |
| **Gene lists** | | | |
| KEGG pathways | Identification of genes coding for metabolic enzymes | https://amp.pharm.mssm.edu/Enrichr/#stats | [35] |
| Extracellular vesicle (EV)-associated genes | Extracellular Vesicle Biogenesis score calculation using GO term IDs GO 0140112, GO 0071971 and GO 0097734 | AmiGO website (http://amigo.geneontology.org/amigo/landing) | |
| | Vesicle-mediated transport score calculation; GO term ID GO 0098876 | | |
| | Vesicle targeting score calculatio using GO term ID GO 0006903 | | |
| | Vesicle fusion score calculation; GO term IDs GO 0006906, GO 0099500 and GO 0031338 | | |
| TCGA RNA-seq dataset | 153 primary GBM | Human Protein Atlas website (https://www.proteinatlas.org/) | |
| TCGA microarray dataset | 485 untreated primary GBM | R2 genomics analysis and visualization platform database (http://r2.amc.nl) | |
| French microarray dataset | 156 GBM (gse 16011) | R2 genomics analysis and visualization platform database (http://r2.amc.nl) | |
| REMBRANDT microarray dataset | 173 GBM | Gliovis portal (http://gliovis.bioinfo.cnio.es) | |
| **Experimental Models** | | | |
| Patient-derived cell ines (PDC) | 6240**, 5706** and R633 PDC obtained from neurosurgical biopsy samples of distinct primary glioblastoma and characterized as descr bed | | [18, 57] |
| Intracranial xenografts | Immunodeficient 8-week old Hsd Athymic Nude-Foxn1nu/Foxn1+ mice | Envigo Laboratories, France | |
| | Stereotaxic injection in the striatum, Coordinates 0 mm posterior and 2.5 mm lateral to the bregma, and 3 mm deep with respect to the surface of the skull | | |
| **Lentiviral vectors** | | | |
| pLKO.1-HPGK-puro-U6-non mammalian shRNA control | control shRNA contruct | Sigma, France | |
| pLKO.1-puro-CMV-tGFP-U6-shELOVL2-64 | ELOVL2 shRNA construct | Sigma, France | |
| **Lentivirus** | control shRNA lentivirus | Plateforme vecteurs viraux et transfert de gènes (Necker Federative structure of research, University Paris Descartes, France) | |
| | ELOVL2 shRNA lentivirus | | |
| **QPCR Primers** | | | |
| ELOVL2 - forward primer | 5'-TCCACTTGGGAAGGAGGCTACA | Eurofins, France | |
| ELOVL2 - reverse primer | 5'-CTCCAAATCAGTAGAGTTCCTGG | Eurofins, France | |
| TBP - forward primer | 5'-TGCACAGGAGCCAAGAGTGAA | Eurofins, France | |
| TBP - reverse primer | 5'-CACATCACAGCTCCCCACCA | Eurofins, France | |
| **Chemicals** | | | |
| Trypan Blue solution 0.4% v/v | 3 min incubation at room temperature; for viable cell counting | Thermo Fisher, France | |
| Puromycin 2 µg/mL | Elimination of non-transduced cells | Sigma, France | |
| Luciferin 20 mM | 150uL intra-peritoneal injection; for bioluminescence imaging | Thermo Fisher, France | |
| **Commercial kits** | | | |
| Nucleospin RNA kit | Total RNA preparation | Macherey-Nagel, France | |
| QuantiTect Reverse Transcription Kit | cDNA preparation | Qiagen, France | |
| SYBR Green PCR Core Reagents kit | QPCR | Bimake.com, France | |
| **Apparatus** | | | |
| Countess automated cell counter | Viable Cell counting | Thermo Fisher, France | |
| LightCycler480 | QPCR | Roche, France | |
| Photon Imager Biospace | Bioluminescence imaging | Biospace Lab, France | |

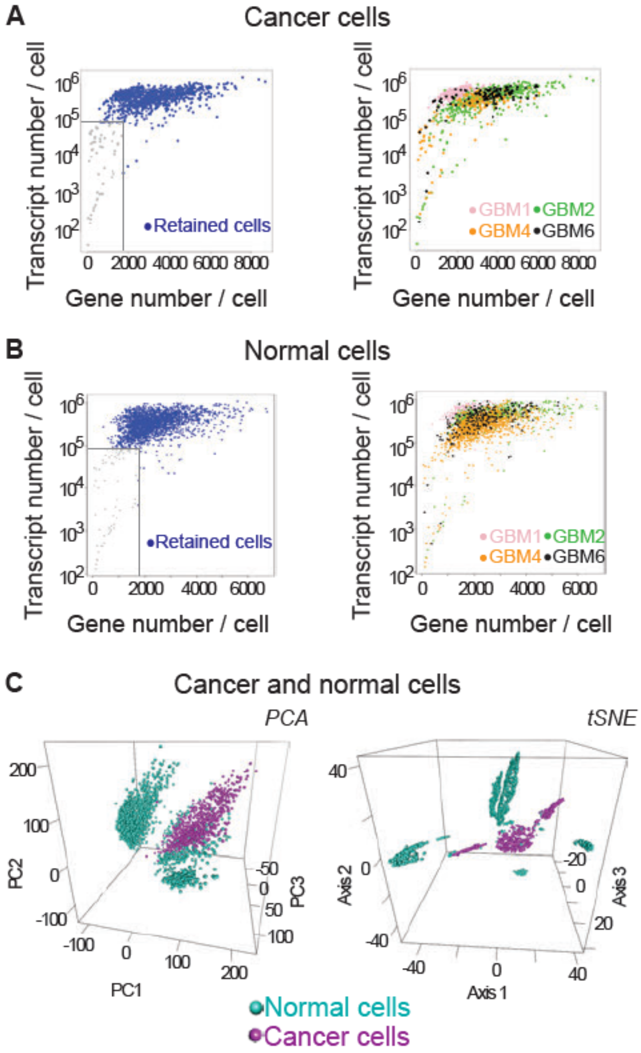


**Fig. S1 (related to Fig.1).**

**Filtering out cells with low-complexity transcriptomes and unsupervised grouping analysis of GBM and normal cells simultaneously.**

**A-B.** Filtering out cells with low-complexity transcriptomes. Cells with more than 90000 transcripts and more than 1700 genes were selected for further analyses. 1033 GBM (A) and 2417 normal cells (B) were retained.
Left panels: selected cells colored in blue, rejected cells colored in gray. Right panels: cells colored by tumor.
**C.** Unsupervised grouping analysis of the mixed set of GBM and normal cells distinguishes cancer cells from normal cells.
Each dot represents a cell. Normal cells colored in light blue, cancer cells colored in violet.
Left panel: PCA visualization. Right panel: tSNE visualization.

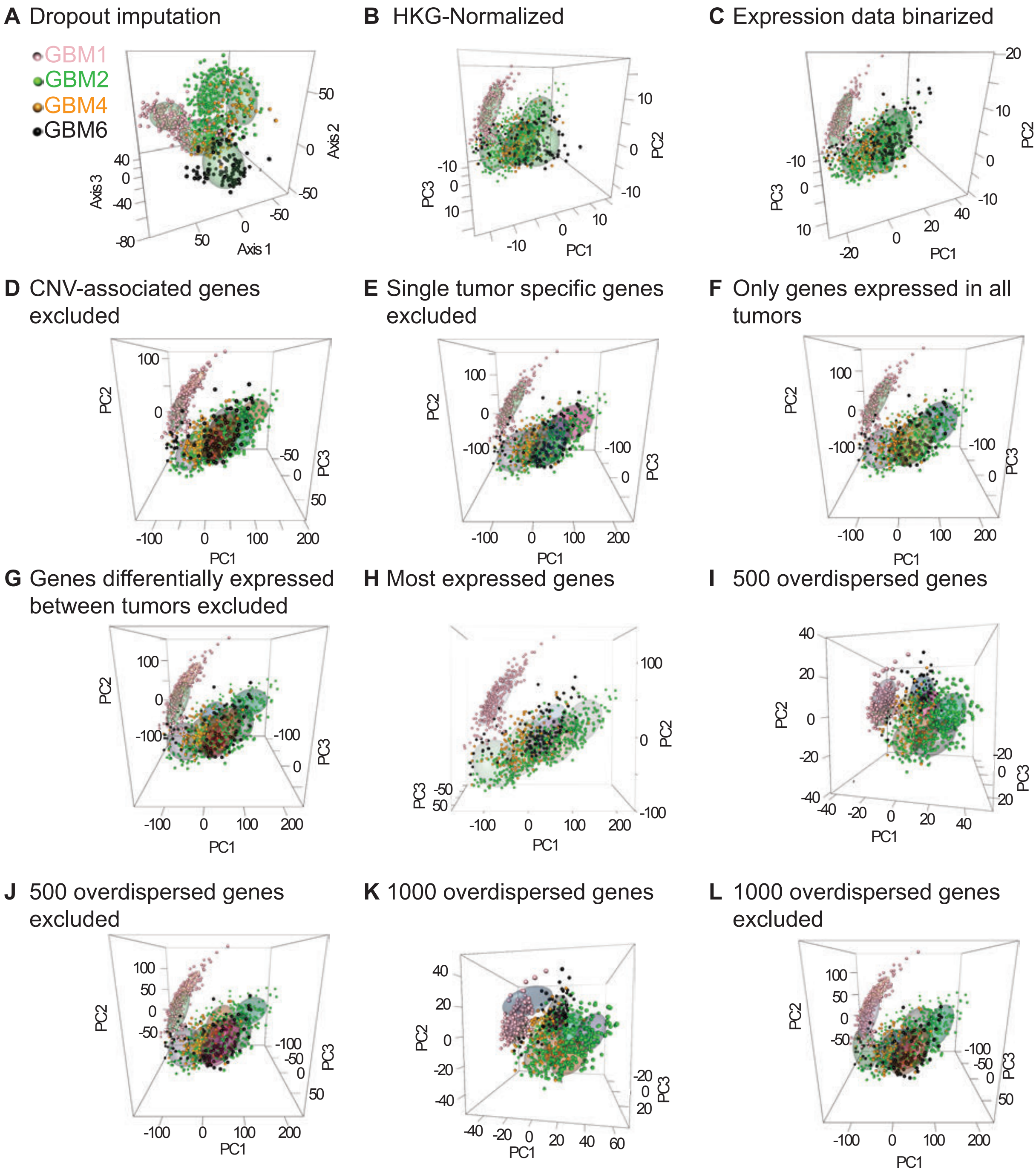


**Fig. S2 (related to Fig. 1).**

**Maintenance of tumor-driven cell grouping regardless of the mode of data normalization or filtering.** A: Multidimensional scaling (MDS) visualization. B-L: PCA visualization. Each dot represents a the clusters identified.

**A-B.** Technical biases related to scRNA-seq do not contribute to tumor-driven cell grouping. A. Maintenance of tumor-driven cell grouping after imputation of dropouts using the algorithm CIDR (Clustering through Imputation and Dimensionality Reduction). CIDR imputes dropouts by inferring their values from gene expression across all cells [42]. **B.** Normalizing data by the expression of housekeeping genes (HKG) prior to analysis fails to alleviate tumor-driven cell grouping.

**C-L.** Tumor-specific biological differences are not reducible to circumscribed sets of genes. **C.** Binarization of expression data to overcome possible inter-experimental variations in the efficacy of scRNA-seq, and therefore in the detected RNA levels, does not change tumor-driven cell grouping. **D.** Filtering out genes located on chromosomes with identified variations in copy number (CNV) does not modify tumor-driven cell grouping. **E.** Tumor-driven cell grouping is maintained after removal of genes detected in a single tumor. **F.** Grouping analysis performed using only genes detected in all tumors also results in tumor-driven cell grouping. **G.** Filtering out genes differentially expressed between tumors does not alleviate tumor-driven cell grouping. **H.** Cell grouping analyses using the most expressed genes across cells fails to overcome tumor-driven cell grouping. **I.** Cell grouping analyses using the top 500 genes with the most overdispersed expression across cells fails to overcome tumor-driven cell grouping. **J.** Filtering out the top 500 genes with the most overdispersed expression across cells does not change tumor-driven cell grouping. **K.** Cell grouping analyses using the top 1000 genes with the most overdispersed expression across cells fails to overcome tumor-driven cell grouping. **L.** Filtering out the top 1000 genes with the most overdispersed expression across cells does not change tumor-driven cell grouping.

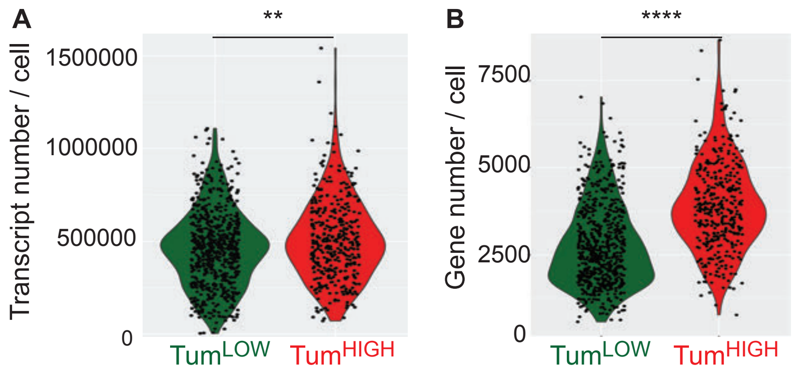


**Fig. S3 (related to Fig. 3)**

**Increased numbers of transcript (A) and genes (B) detected per cell with high tumorigenic scores, compared to cells with low tumorigenic scores.**
Mann-Whitney test. p = 1.92*10-3 for transcript number
and 2*10-41 for gene number.

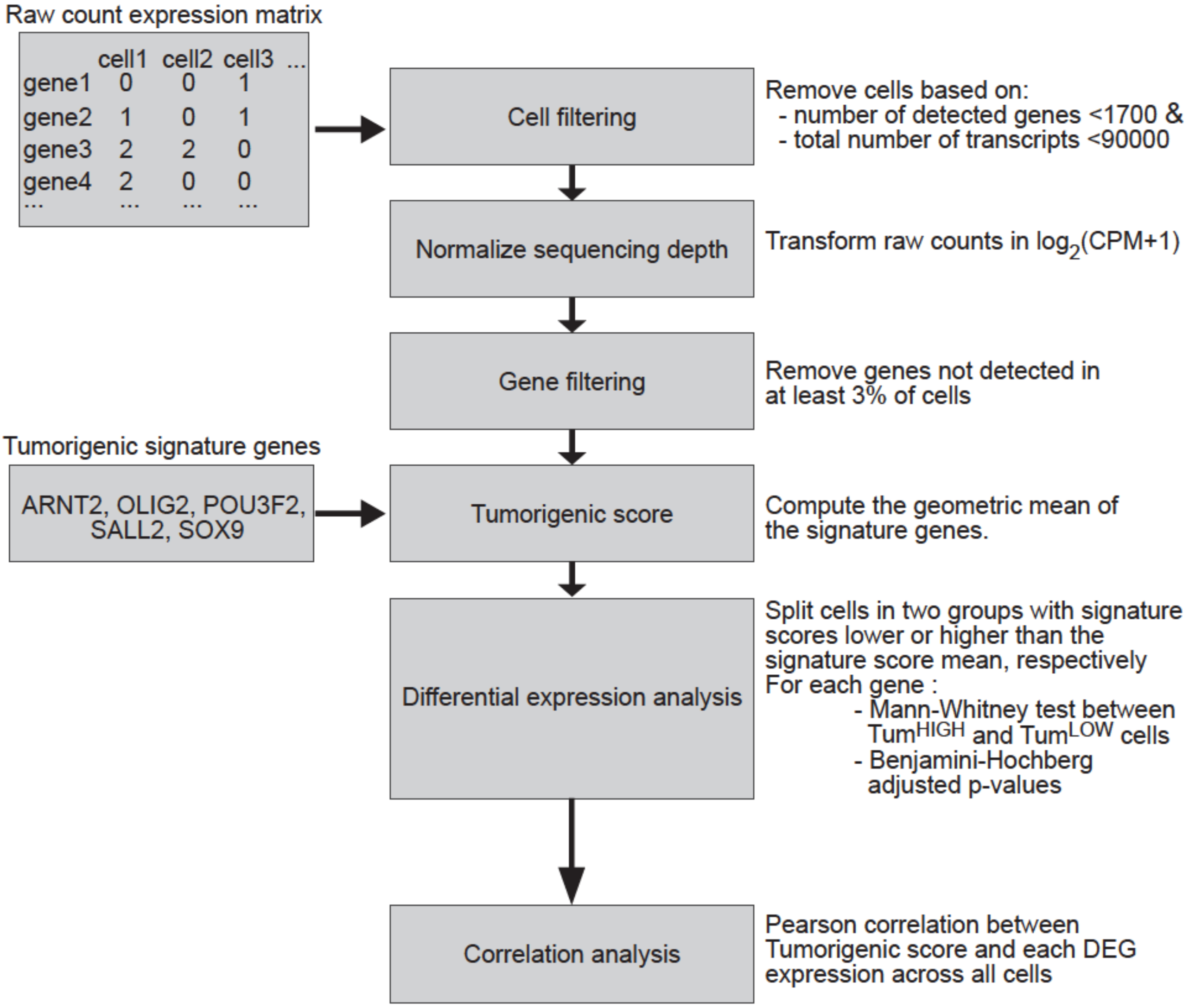


**Fig. S4. Signature-based analytical workflow.**

The analytical method developed has been implemented in R (Datafile S2).

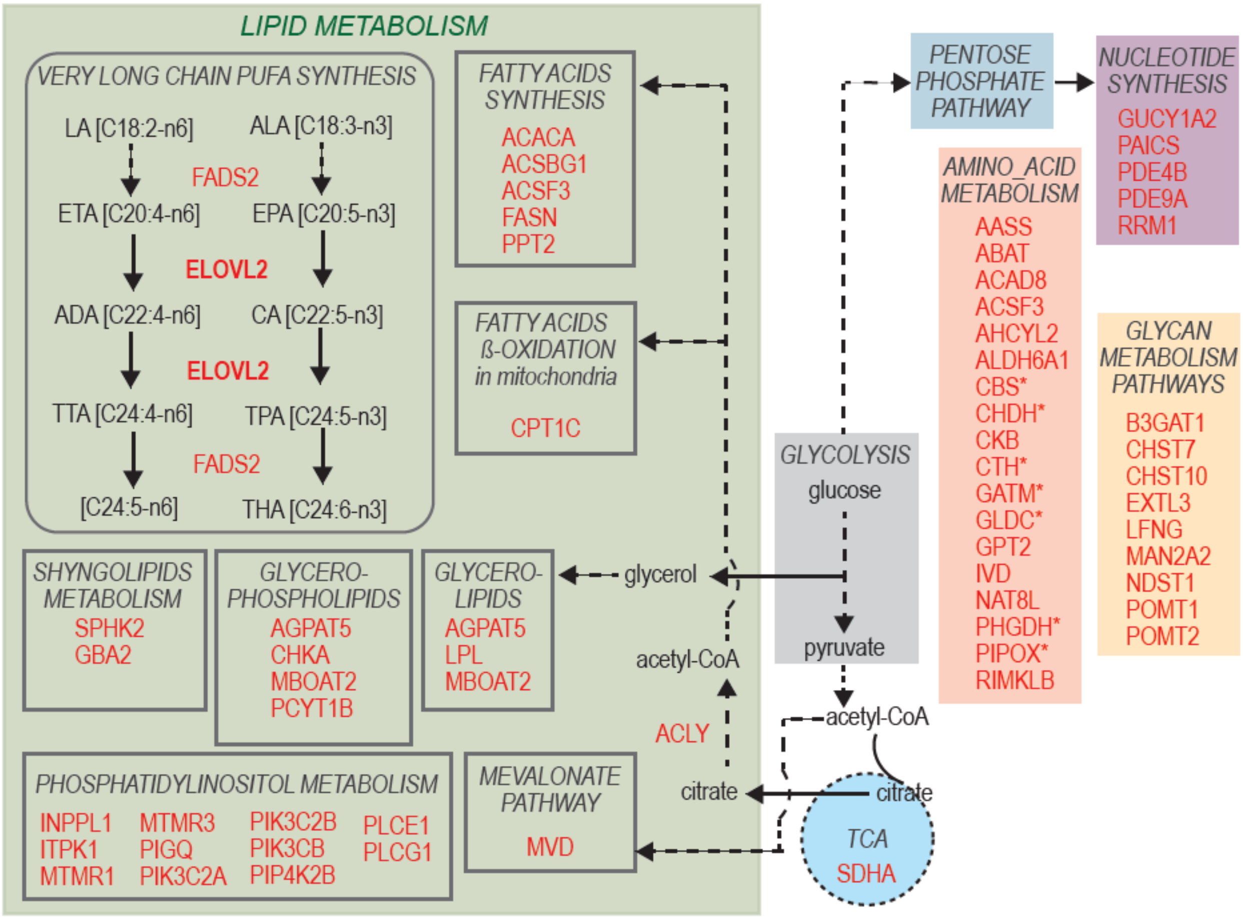


**Fig. S5 (related to Fig. 4)**

**Schematic representation of the main metabolic pathways in which are involved 60 of the 66 metabolic enzyme genes overexpressed in TumHIGH GBM cells and tissues.**

Asterisks mark genes coding for components of the glycine, serine and threonine metabolism.

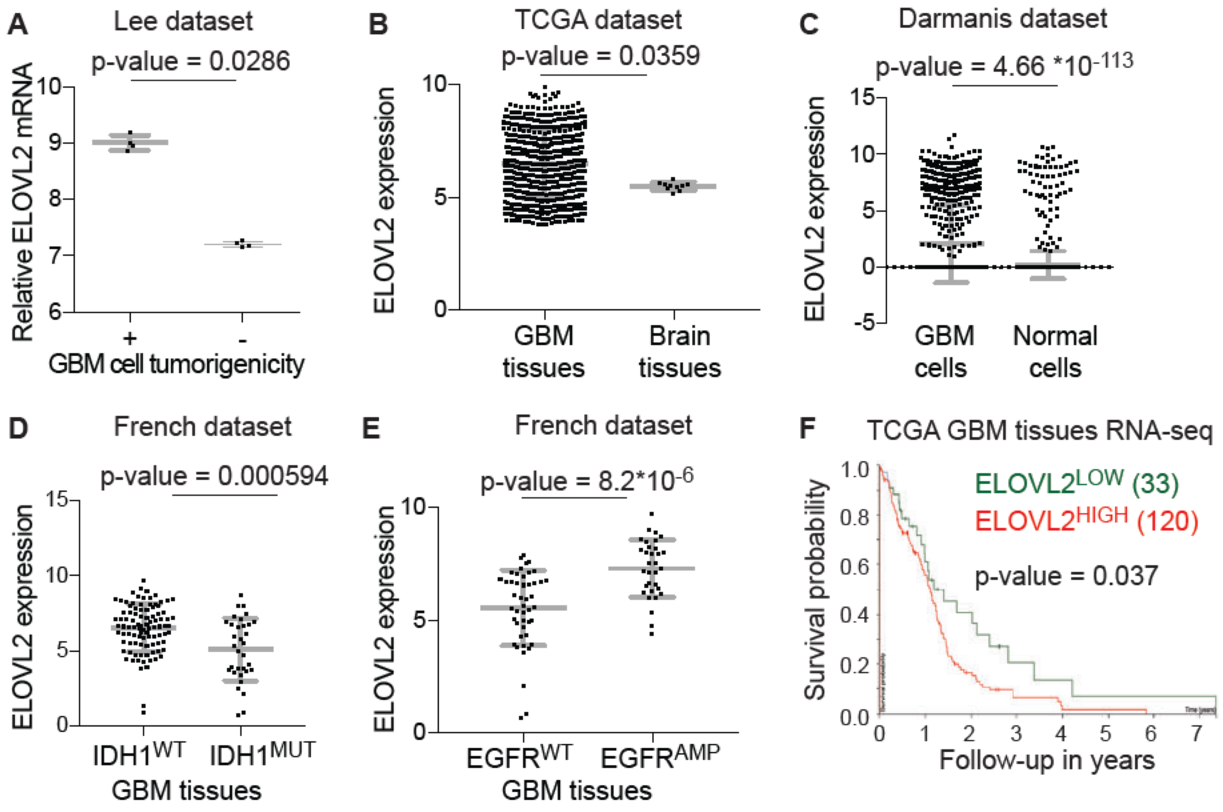


**Fig. S6 (related to Fig. 5)**

**Increased ELOVL2 expression is associated with increased tumor burden.**

**A.** Down-regulated ELOVL2 expression in patient-derived GBM cells deprived of tumorigenic properties, compared to their tumorigenic counterparts. Mann-Whitney test. Lee microarray dataset GEO ID: GSE4536. R2 genomics analysis and visualization platform database (http://r2.amc.nl).

**B.** Higher ELOVL2 expression in GBM tissues compared to normal brain tissues. Mann-Whitney test. TCGA tissue transcriptome dataset (microarrays) of 528 primary GBM and 10 normal brain tissues.

**C.** Significantly higher ELOVL2 expression in GBM cells compared to normal cells. Mann-Whitney test. scRNA-seq dataset from Darmanis and colleagues [14].

**D-E.** ELOVL2 expression prevails in GBM bearing a wild-type form of IDH1 (D) and with EGFR gene amplification (EGFRAMP) (E). French tissue transcriptome dataset (microarrays). IDH1WT, n=95. IDH1MUT n=33. EGFRWT, n=46. EGFRAMP, n=32. Mann-Whitney test. F. High ELOVL2 expression is associated with a poorer survival for patients. TCGA GBM tissue transcriptomes (RNA-seq) of 153 primary GBM. Log rank test.

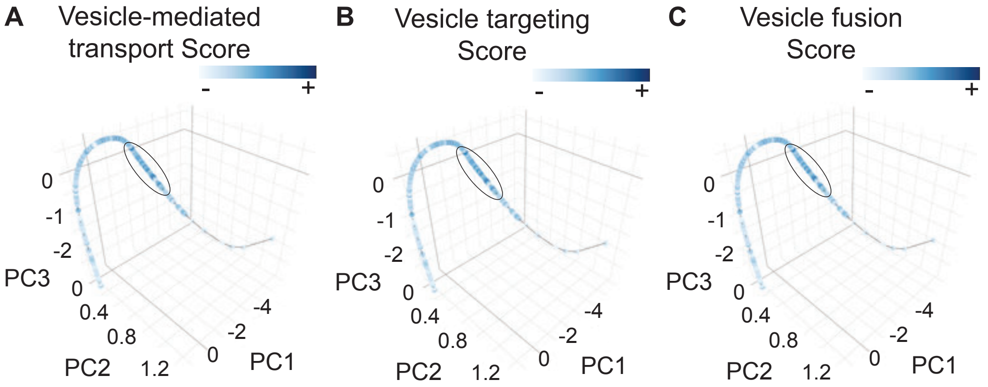


**Fig. S7 (related to Fig. 5)**

**Principal curve analysis associates GBM cell tumorigenic state to mobilization of vesicle production and release.**
Principal curve resulting from PCA of the expression of the subgroup of genes encoding lipid metabolism enzymes overexpressed in TumHIGH cells and tissues. Cells colored according to their score calculated with the components of molecular signatures associated with extracellular vesicle transport (A), targeting (B), and fusion (C). Note that cells with either high score cluster on the same portion of the curve (ellipses).
